# LRG1 destabilizes tumor vessels and restricts immunotherapeutic potency

**DOI:** 10.1101/2020.10.12.334359

**Authors:** Marie N. O’Connor, David M. Kallenberg, Rene Jackstadt, Angharad H. Watson, Markella Alatsatianos, Julia Ohme, Carlotta Camilli, Camilla Pilotti, Athina Dritsoula, Chantelle E. Bowers, Laura Dowsett, Jestin George, Xiaomeng Wang, Ann Ager, Owen J. Sansom, Stephen E. Moss, John Greenwood

**Author notes:** Joint contributing authors (Marie N. O’Connor, David M. Kallenberg). Joint last and corresponding authors.

## Abstract

Vascular dysfunction contributes to the pro-oncogenic tumor microenvironment and impedes the delivery of therapeutics. Normalizing of the tumor vasculature has therefore become a potential therapeutic objective. We previously reported that the secreted glycoprotein, leucine-rich α-2-glycoprotein 1 (LRG1), contributes to the formation of pathogenic neovascularization. Here we show that in mouse models of cancer, *Lrg1* is induced in tumor endothelial cells. We demonstrate that the expression of LRG1 impacts on tumor progression as *Lrg1* deletion or treatment with a LRG1 function-blocking antibody inhibited tumor growth and improved survival. Inhibition of LRG1 increased endothelial cell pericyte coverage and improved vascular function resulting in significantly enhanced efficacy of cisplatin chemotherapy, adoptive T-cell therapy and immune checkpoint inhibition (anti-PD1) therapy. With immunotherapy, LRG1 inhibition led to a significant shift in the tumor microenvironment from being predominantly immune silent (cold) to immune active (hot). LRG1 therefore drives vascular abnormalization and its inhibition represents a novel and effective means of improving the efficacy of cancer therapeutics.

## INTRODUCTION

Angiogenesis is notably different in development and disease, with the former producing an organized, stable and functional vascular network, and the latter being typically disorganized and dysfunctional. Yet vascularization in both settings is driven by many of the same molecules, such as vascular endothelial growth factor (VEGF). Why vessels fail to grow in a patterned and functional manner in most disease settings remains poorly understood but points to the potential involvement of novel pathogenic factors that corrupt physiological angiogenesis. We and others have previously shown that in vascular pathology of the retina (*1*) and kidney (*2*) the secreted glycoprotein LRG1 is induced and promotes dysfunctional vessel growth through modifying endothelial cell TGFβ signaling. Importantly, deletion of *Lrg1* did not impact on developmental angiogenesis with mice exhibiting no overt phenotype (*1*). These observations suggest that in disease LRG1 might be a contributing pathogenic factor responsible for preventing the development of physiological vessels and thus play a role in the vascular dysfunction that is prevalent in cancer.

The formation of new blood vessels has long been recognized as an essential feature of tumor expansion, survival and metastatic spread (*3,4*). Consequently, targeting of key pro-angiogenic signaling molecules, most notably VEGF through blocking antibodies such as bevacizumab, to limit vascular growth or to regress existing vessels has become an established therapeutic regimen. Whilst such approaches have met with some success, resulting in an increase in progression-free survival in certain cancers, they have had little impact on overall survival rate. As in other diseases, cardinal features of tumor vessels are that they are poorly perfused, leaky and hemorrhagic; characteristics believed to be due in part to the failure of vessel maturation. These abnormal features result in an hypoxic pro-oncogenic tumor microenvironment (TME) that promotes malignancy and metastatic spread, impairs beneficial immune responses, and limits the efficacy of systemically administered drugs and immunotherapeutics (*5,6*). Counter to the original rationale of blocking neovascularization, therefore, an alternative strategy has emerged which aims to normalize the vasculature and render the microenvironment more conducive to tumor destruction (*7,8*). In pursuit of this objective, various approaches have been tested including the use of anti-VEGF drugs delivered at a lower dose than that required to prevent, or ablate, neovascularization (*9*). This tactic has led to some improvement in the delivery of cancer therapeutics (*10,11*) but the timing and dosage remain problematic, especially as the window of opportunity with anti-VEGF drugs may be transient (*10,12,13*). To overcome this limitation other vascular modifying approaches have been investigated, either as monotherapy or in conjunction with existing anti-VEGF drugs (*14–22*). These studies have demonstrated that sustained vascular normalization can be achieved, at least in the experimental setting, and validate the potential utility of these strategies in enhancing the efficacy of current standard of care and emerging treatments. Moreover, in the context of vascular normalization and immunotherapy, these studies also highlight the importance of crosstalk between the vasculature and the immune system in establishing a favorable therapeutic milieu (*19,23–26*). In particular, it has been shown that vascular normalization strategies combined with checkpoint inhibition result in the formation of high endothelial venule (HEV) characteristics within the tumor vasculature that help promote the recruitment of effector T cells (*14*). It is clear, therefore, that there is much that we still need to understand about the contribution that the vasculature makes to tumor progression and in so doing reveal new potential therapeutic targets.

The discovery that LRG1 is associated with abnormal vessel growth in various diseases (*1,2*) raises the possibility that this secreted glycoprotein is a contributing factor in abnormal tumor vessel growth. Consistent with this hypothesis, studies have shown *LRG1* to be induced in many carcinomas, and there is growing evidence that raised blood LRG1 levels alone, or in combination with other biomarkers, correlate with increased tumor load and poor prognosis (Supplementary Table 1). Such data strongly implicate LRG1 in the pathogenesis of cancer and provide a rationale for further investigation. In this study, therefore, we aimed to establish whether LRG1 impacts on tumor vessel structure and function and crucially the implications of this on therapy. We show that LRG1 affects vessel growth, structure and function establishing it as a significant vascular destabilizing factor. By deleting *Lrg1*, or blocking its function with a targeted antibody, we observe reduced tumor growth, and demonstrate that tumor vessels exhibit a more physiological configuration with improved pericyte coverage. The ramifications of vessel normalization brought about by LRG1 blockade were further investigated and revealed enhanced efficacy when combined with cytotoxic and immunotherapeutic strategies. These data provide evidence that blockade of LRG1 in cancer offers a novel approach to vascular normalization and in doing so, potentiates the efficacy of current and emerging therapies.

## RESULTS

### *Lrg1* deletion reduces tumor growth and increases survival

To test the hypothesis that LRG1 contributes to tumor growth via vascular destabilization, we first examined *Lrg1* expression in the syngeneic B16-F0 mouse melanoma and Lewis lung carcinoma (LLC) subcutaneous graft models, the KPC model of pancreatic ductal adenocarcinoma (PDAC) (*LSL-Kras^G12D/+^; LSL-Trp53^R172H/+^*; *Pdx-1-Cre*), the *Apc^Min/+^* and the *vil^CreER^ Apc^fl/+^* models of colorectal cancer (CRC). *Lrg1* transcript was detected within the tumors of all models (Fig. 1A-C). In B16-F0 and LLC tumors, *Lrg1* expression appeared to be restricted mostly to vessels (Fig. 1A) whereas in intestinal adenomas and PDAC, *Lrg1* expression was highly upregulated in the neoplasm compared to normal tissue (Fig. 1B,C. Supplementary Fig. 1). Consistent throughout the different cancer models *Lrg1*, which is not detectable in normal endothelial cells, was induced but was not observed in αSMA positive perivascular mural cells (Fig. 1D). Overall, these observations corroborate reports in human cancers (Supplementary Table 1) and raise the possibility that the secretion of LRG1 may impact on the tumor microenvironment and, through both autocrine and paracrine signaling, on vascular function.

**Figure 1.**
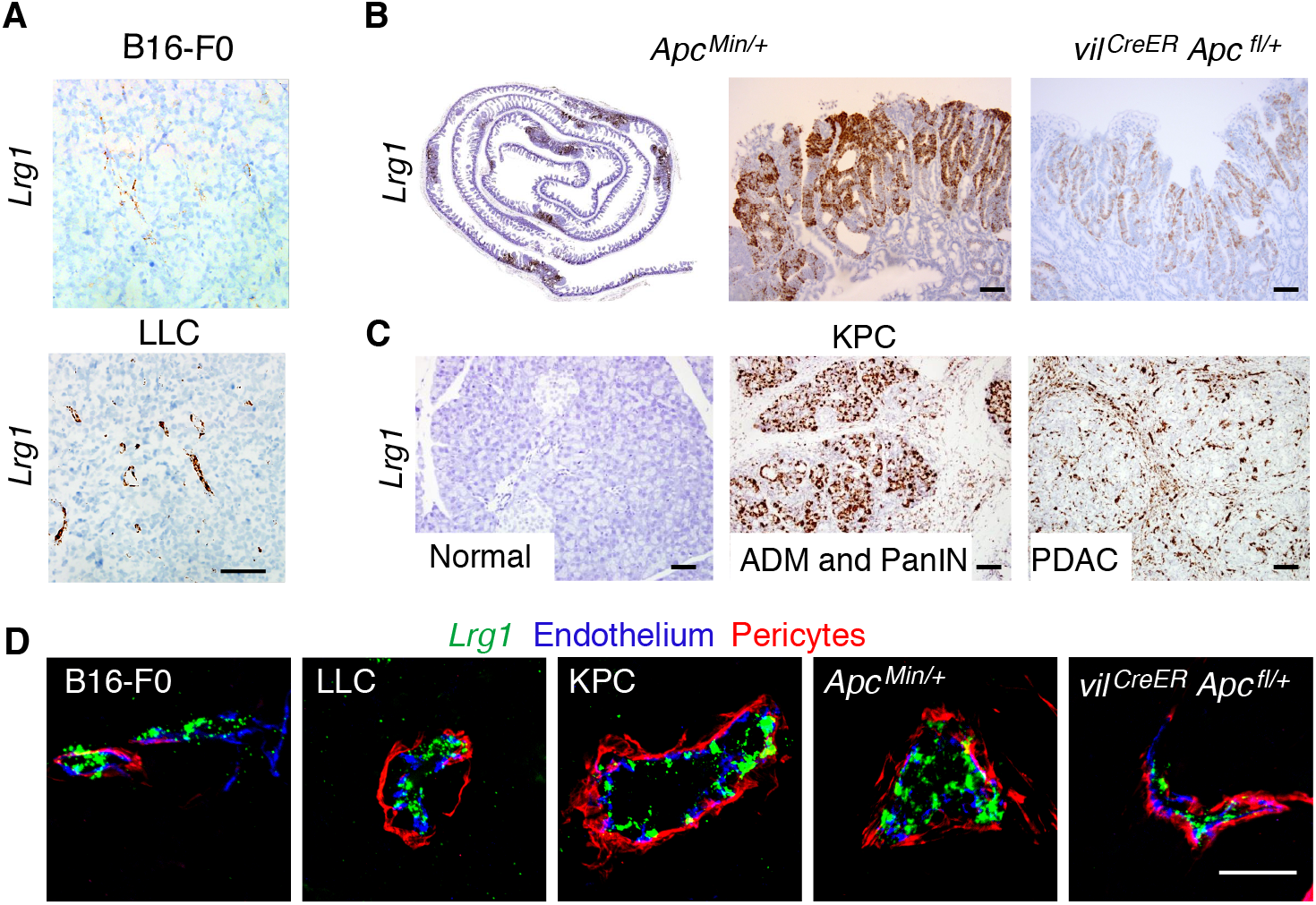
*Lrg1* gene is induced in tumor endothelial cells and in some cancers. *Lrg1* transcript expression in (**A**) B16-F0 and LLC syngeneic tumors, (**B**), *Apc^Min/+^* and *vil^CreER^ Apc^fl/+^* small intestine and (**C**), normal or diseased pancreas showing acinar ductal carcinoma (ADM), pancreatic intraepithelial neoplasia (PanIN) or PDAC from KPC mice. Scale bar 50μm. (**D**). Vascular *Lrg1* transcript expression (green) and immunohistochemical cell markers for EC (CD31; blue) and pericytes (αSMA; red). Scale bar, 30 μm.

Our previous work demonstrated that *Lrg1* knockout, or functional blockade with an antibody, reduced the size of neovascular lesions in models of age-related macular degeneration and ocular hypoxia. We therefore examined the effects of *Lrg1* gene deletion on tumor growth and survival across a range of syngeneic and genetic models. In the B16-F0 and LLC subcutaneous tumors, tumor growth was significantly reduced in *Lrg1*^−/−^ mice compared to WT controls (Fig. 2A,B. Supplementary Fig. 2), with a decrease in final tumor volume at the termination end-point of 44% and 46%, respectively. Consistent with these observations, both the *Apc^Min/+^* (Fig. 2C) and *vil^CreER^ Apc^fl/+^* (Fig. 1D) mice exhibited a significantly enhanced survival rate on the *Lrg1*^−/−^ background. In both colorectal models there was a trend towards a reduced tumor number in *Lrg1*^−/−^ mice that reached significance in the colon and small intestine in the *Apc^Min/+^* and *vil^CreER^ Apc^fl/+^* mice respectively (Fig. 2E,F). We next investigated whether knockout of *Lrg1* affected the survival of the KPC pancreatic ductal adenocarcinoma (PDAC) tumor bearing mice (*27*). As with the other models, *Lrg1*^−/−^ KPC mice exhibited a significantly enhanced survival rate compared to *Lrg1*^+/+^ mice (Fig. 2G). These results demonstrate that deletion of *Lrg1*^−/−^ in a number of different tumor models improves outcome and supports the hypothesis that LRG1 is pro-oncogenic.

**Figure 2.**
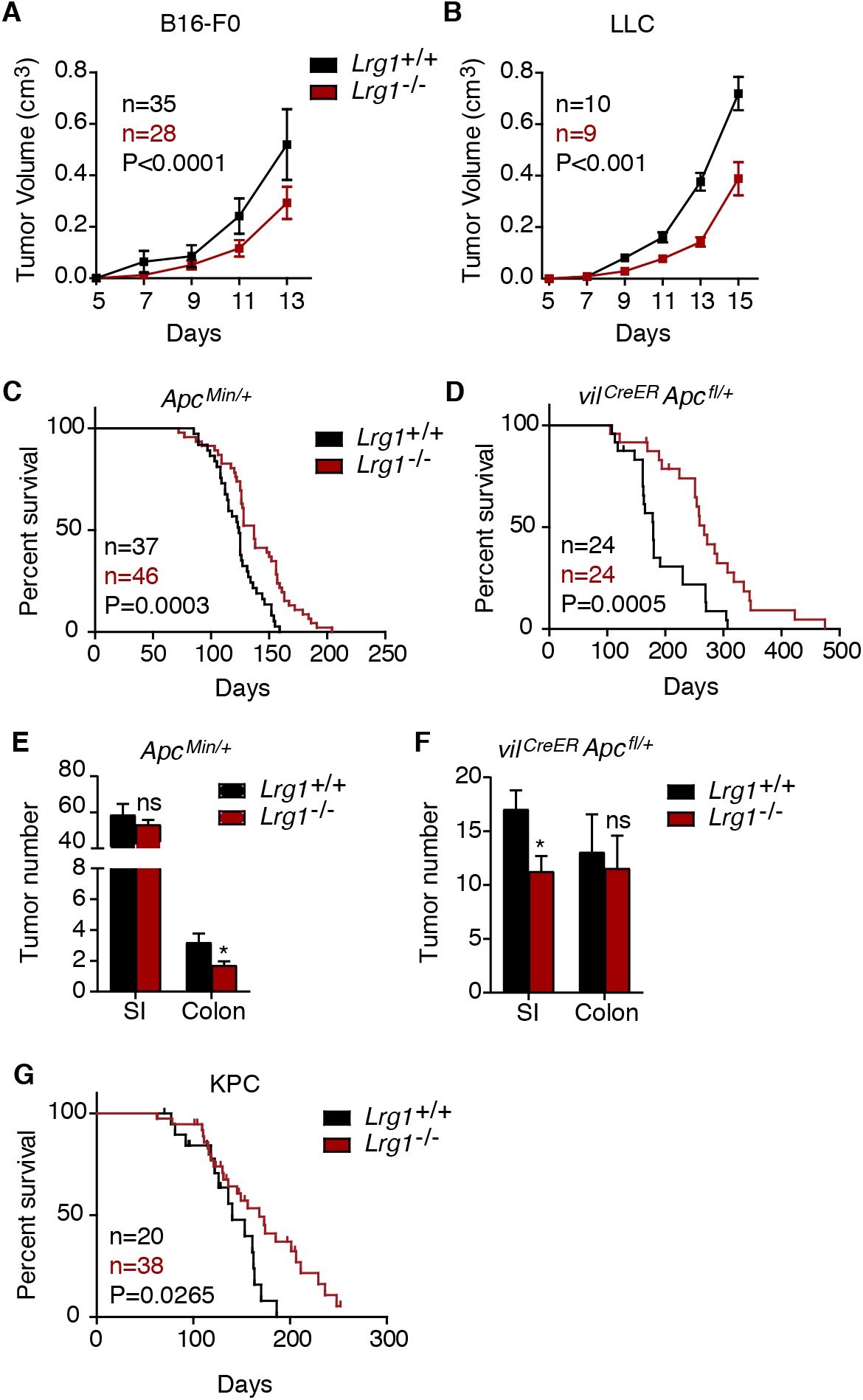
*Lrg1* deletion reduces tumor volume and enhances survival. Growth curves of (**A**) B16-F0 and (**B**) LLC subcutaneous tumors from *Lrg1*^+/+^ or *Lrg1*^−/−^ mice (mean ± 95% CI). RM two-way ANOVA. (**C**), Kaplan-Meier survival curve of *Apc^Min/+^* and (**D**), *vil^CreER^ Apc^fl/+^* mice with or without homozygous deletion of *Lrg1*. Mantel-Cox test. (**E**). Tumor number in *Apc^Min/+^* (*n*=16 *Lrg1^+/+^* and *n*=27 *Lrg1*^−/−^) and (**F**), *vil^CreER^ Apc^fl/+^* (*n*=11 *Lrg1^+/+^* and *n*=11 *Lrg1*^−/−^) mice with or without homozygous deletion of *Lrg1*. (**G**), Kaplan-Meier survival curve of KPC mice with or without homozygous deletion of *Lrg1*. Mantel-Cox test.

### *Lrg1* deletion results in tumor vessel normalization

To ascertain if reduced tumor size and enhanced survival in *Lrg1*^−/−^ mice correlated with changes in vascularization we measured the percentage of vessel area in each of the tumors. No difference in total vessel area between wild type and *Lrg1* knockout mice was observed across the different models (Fig. 3A,B. Supplementary Fig. 3A). There was only a modest reduction in the number of vessel profiles per unit area in the B16-F0, LLC and KPC mice on the *Lrg1*^−/−^ background, but a striking increase in the size of individual vessels (Fig. 3B,C). Due to the planar orientation of the tumor vasculature in the *Apc^Min/+^* models, comparable analysis of vessel density was not possible. Nonetheless, our observations that endothelial cells express the *Lrg1* gene in a pathological setting and that tumor vessels tend to be larger, raise the possibility that LRG1 impacts tumor vascularization.

**Figure 3.**
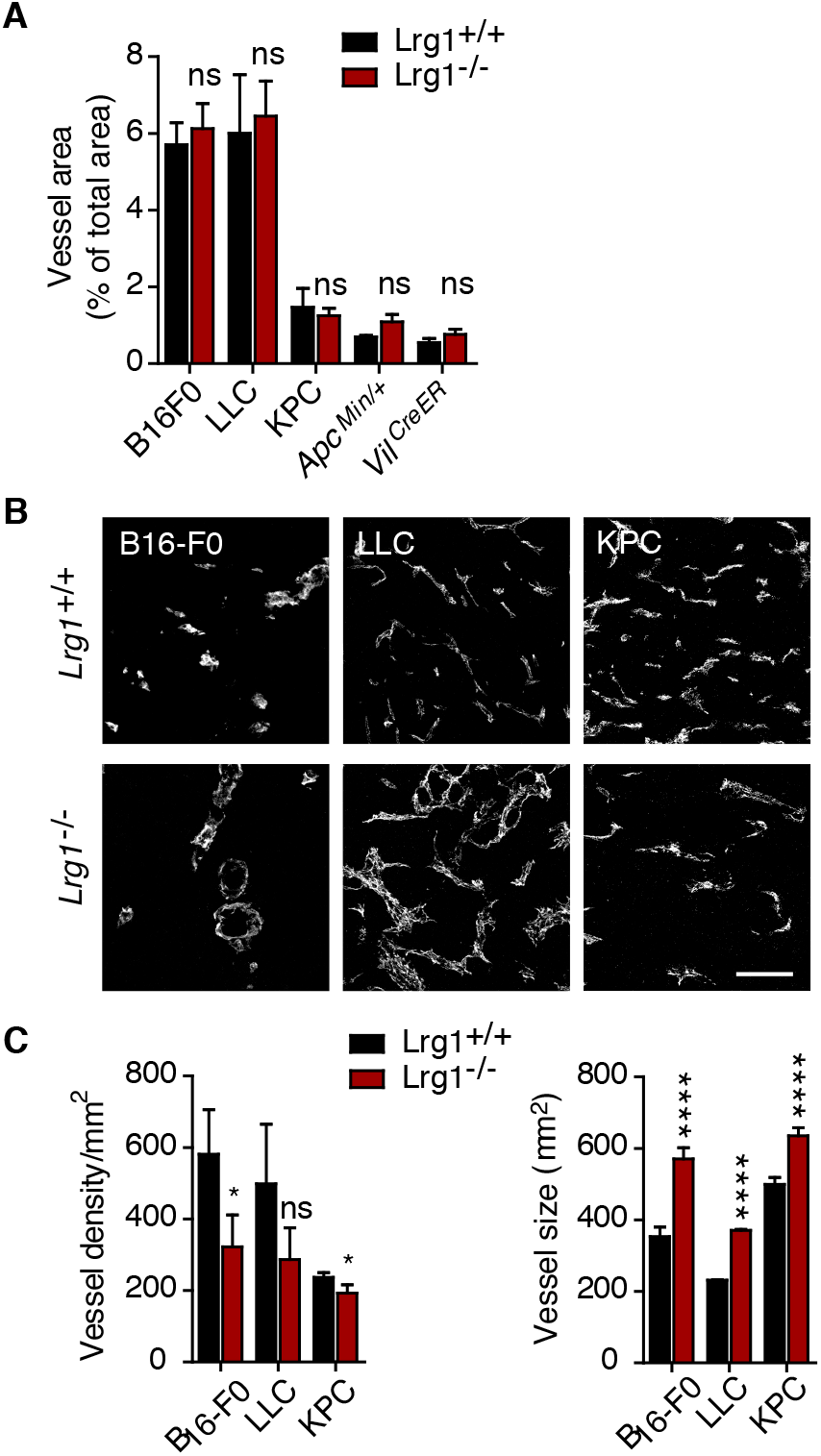
*Lrg1* deletion impacts on vascular structure. Vessel area (**A**) for different tumors expressed as the percentage of field that was positive for the endothelial cell marker CD31. (**B**). CD31 stained sections of the vasculature from B16-F0, LLC and KPC tumors from *Lrg1*^+/+^ and *Lrg1*^−/−^ mice (scale bar, 50 μm) and (**C**) quantification of vessel density and size (cross-sectional area) of individual CD31^+^ vessels. B16-F0 tumors (*n*=15 *Lrg1^+/+^* and *n*=17 *Lrg1*^−/−^), LLC tumors (*n*=28 *Lrg1^+/+^* and *n*=25 *Lrg1*^−/−^), KPC tumors (*n*=7 *Lrg1^+/+^* and *n*=10 *Lrg1*^−/−^), *Apc*^Min/+^ tumors (*n*=8 *Lrg1^+/+^* and *n*=15 *Lrg1*^−/−^) and vil^CreER^ *Apc*^fl/+^ tumors (*n*=8 *Lrg1^+/+^* and *n*=8 *Lrg1*^−/−^). (**H**-**L**) Mean ± 95% CI. Mann Whitney test, *P<0.05, ****P<0.0001, ns non-significant.

The observed loss of small vessels, with a concomitant increase in larger vessels, is a characteristic associated with vessel normalization (*13*) and suggests that the presence of LRG1 may impair vessel maturation. Pericyte coverage and basement membrane deposition are additional features associated with vessel stabilization and maturation (28), and failure of these processes is known to contribute to tumor vessel dysfunction (29). We therefore investigated whether the absence of LRG1 affects this critical relationship. Using NG2 and/or αSMA as markers of mural cells, we observed that *Lrg1* deletion in the B16-F0, *Apc^Min/+^* and *vil^CreER^ Apc^fl/+^* models resulted in an increase in mural cell coverage of endothelial cells (Fig. 4A,B). Whilst there was also an increase in coverage in the LLC and the KPC models, this did not reach significance. Similar to mural cell coverage, vessels from B16-F0, *Apc^Min/+^* and *vil^CreER^ Apc^fl/+^* tumors, but not those from the LLC or KPC tumors, also exhibited increased basement membrane association (Fig. 4C; Supplementary Fig. 3B). In further support of the observation that LRG1 depletion improves vascular structure, scanning electron micrographs of B16-F0 tumors from *Lrg1*^−/−^ mice revealed a reduction in the amount of intraluminal membranous inclusions (Fig. 4D), a recognized feature of abnormal tumor vessels (*30*). These data indicate that not only does *Lrg1* knockout impact on tumor size and survival, but its loss is also associated with the acquisition of a more normal vascular appearance.

**Figure 4.**
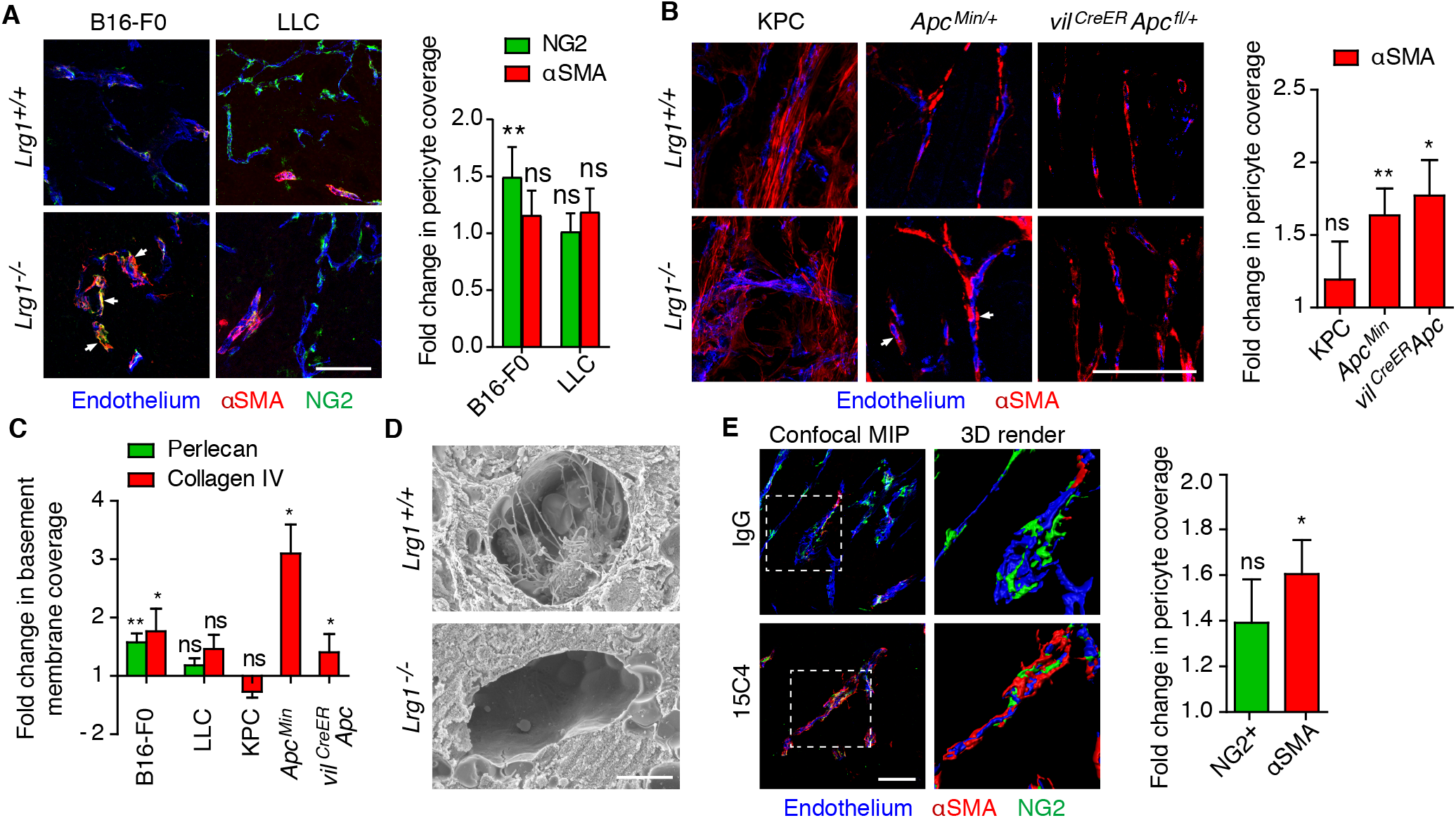
Loss or inhibition of LRG1 improves blood vessel structure. Pericyte (NG2 or αSMA) association with tumor EC (CD31 or podocalyxin) in the B16-F0, LLC, KPC, *Apc^Min/+^* and *vil^CreER^ Apc^fl/+^* tumor models with or without homozygous deletion of *Lrg1* (**A**,**B**). Tight association of NG2^+^ pericytes with the tumor vessel is indicated by arrowheads (**A**). KPC and *Apc^Min/+^* sections were labelled with antibodies to CD31 and αSMA (**B**). For NG2, B16-F0 (*n*=10 *Lrg1^+/+^* and *n*=15 *Lrg1*^−/−^ mice) and LLC (*n*=14 *Lrg1^+/+^* and *n*=15 *Lrg1*^−/−^ mice). For αSMA, B16-F0 (*n*=11 *Lrg1^+/+^* and *n*=15 *Lrg1*^−/−^ mice), LLC (*n*=7 *Lrg1^+/+^* and *n*=7 *Lrg1*^−/−^ mice), KPC (*n*=5 *Lrg1^+/+^* and *n*=10 *Lrg1*^−/−^ mice) and *Apc*^Min/+^ (mean values from *n*=8 *Lrg1^+/+^* and *n*=11 *Lrg1*^−/−^ mice); Scale bars, 100 μm. Student t-test. (**C**), Endothelial basement membrane (perlecan and/or collagen IV) association with tumor endothelium (CD31). For perlecan, B16-F0 (*n*=12 *Lrg1^+/+^* and *n*=11 *Lrg1*^−/−^ mice) and LLC (*n*=19 *Lrg1^+/+^* and *n*=18 *Lrg1*^−/−^ mice). For collagen IV, B16-F0 (*n*=12 *Lrg1^+/+^* and *n*=11 *Lrg1*^−/−^ mice), LLC (*n*=19 *Lrg1^+/+^* and *n*=18 *Lrg1*^−/−^ mice), KPC (*n*=4 *Lrg1^+/+^* and *n*=8 *Lrg1*^−/−^ mice) and *Apc*^Min/+^ (mean per mouse, *n*=5 *Lrg1^+/+^* and *n*=10 *Lrg1*^−/−^ mice); Mann Whitney, ns non-significant, *P<0.05, **P<0.01. Mann Whitney. (**D**), Scanning electron microscopy of B16-F0 tumors grown in *Lrg1*^+/+^ or *Lrg1*^−/−^ mice. Scale bar, 5 μm. (**E**), Immunohistochemistry and quantification of pericyte (NG2 and αSMA) association with B16-F0 tumor endothelium (CD31 or podocalyxin) from wild-type mice treated with anti-LRG1 (15C4) or control antibody (IgG). Scale bar, 100 μm. 3D renders of the highlighted areas are shown. Graph shows fold change (mean ± s.e.m.) in pericyte overlap. For NG2, IgG *n*=11, 15C4 *n*=12. For αSMA, IgG *n*=16, 15C4 *n*=19 tumors; Student t-test, ns non-significant; Student t-test. *P<0.05.

To investigate whether *Lrg1* knockout alters gene expression of key signaling axes genes involved in either vascular maturation or destabilization we undertook RNASeq analysis. Accordingly, we investigated signature genes for the receptor-ligand pathways of VEGF, angiopoietin (ANGPT), platelet derived growth factor (PDGF), TGFβ, sphingosine 1-phosphate (S1P), Notch, Wnt, Hedgehog, fibroblast growth factor, ephrin and apelin (Supplementary Fig. 4). None of these signature genes showed evidence of significant alteration indicating that their dysregulation is not responsible for the LRG1 effects, that LRG1 may operate post-translationally by impacting on signaling or that it mediates its effect by alternative mechanisms. We also investigated the expression of key glycolytic pathway genes that have been associated with angiogenesis (*31*) and vascular dysregulation (*32*). No significant differences were observed except for an increase in the flow-sensitive transcription factor Krüppel-like factor 2 (*Klf2*), which has been reported to repress endothelial cell metabolism via PFKFB3 (*33*) and impact on angiogenesis through this and other angiogenic pathways. Up-regulation of *Klf2* is consistent with a shift from turbulent oscillatory flow to uniform laminar flow and is in accordance with the observed vascular normalization (*34*).

We also investigated the gene expression profiles of key endothelial cell adhesion molecules as these have been reported to be suppressed in tumor vasculature and contribute to endothelial cell anergy. We observed elevated expression of most of the common adhesion molecules, with *Icam1* and *Vcam1* being significantly increased (Supplementary Fig. 5), further supporting the contention that the vasculature is normalized by *Lrg1* deletion.

### LRG1 antibody blockade results in improved vascular function

Having demonstrated that deletion of *Lrg1* influences tumor growth and vascular organization, we next asked whether this could be phenocopied by inhibiting the activity of LRG1 in wild type mice with a LRG1 function-blocking antibody. For this we chose the B16-F0 tumor-bearing mice which exhibited robust effects of *Lrg1* deletion and which are generally considered to respond poorly to therapeutic intervention. Following a subcutaneous graft of B16-F0 cells, mice were treated intraperitoneally with 15C4, a LRG1 function-blocking monoclonal antibody (35). As with *Lrg1*^−/−^ mice, LRG1 antibody blockade resulted in a similar reduction in tumor volume of 39% at the experimental endpoint (Supplementary Fig. 6A,B), a decrease in vessel density, an increase in vessel size (Supplementary Fig 6C), and an increase in mural cell association with tumor vascular endothelial cells (Fig. 4E). However, the increase in basement membrane coverage was not significant (Supplementary Fig. 6D).

These data show that inhibition of LRG1 results in a more physiological vascular configuration that might be expected to be associated with vessel stabilization and improved vascular function. To test this assertion, we examined whether inhibition of LRG1 leads to an increase in tumor vessel perfusion. Using a systemically delivered fluorescent lectin tracer to mark perfused vessels, and an antibody to decorate all endothelial cells in tissue sections, we observed a significant increase in the percentage of perfused tumor vessels in mice treated with 15C4 (Fig. 5A). We then asked whether this increase in vascular patency reduced tumor hypoxia, and found that treatment of B16-F0 bearing mice with 15C4 indeed led to a significant reduction in tumor hypoxia (Fig. 5B). Another feature of vascular normalization is reduced vascular permeability through the stabilization of endothelial cell junctions. The adherens junction protein VE-cadherin regulates vascular endothelial cell junction integrity and its enhanced expression is associated with a reduction in vascular leakage (*36*). In 15C4 treated mice we observed a significant increase in the intensity of staining of endothelial VE-cadherin (Fig 5C) consistent with improved barrier integrity. In addition, we saw a reduction in tumor vessel permeability as indicated by less diffusion of Hoechst dye from perfused (lectin positive) vessels (Fig. 5D). The vascular normalization also resulted in a small but non-significant increase in CD3 positive T cells (Fig 5E). Taken together, these data indicate that LRG1 blockade improves vascular function and in so doing confirms LRG1 both as an angiopathic factor and potential therapeutic target in tumors.

**Figure 5.**
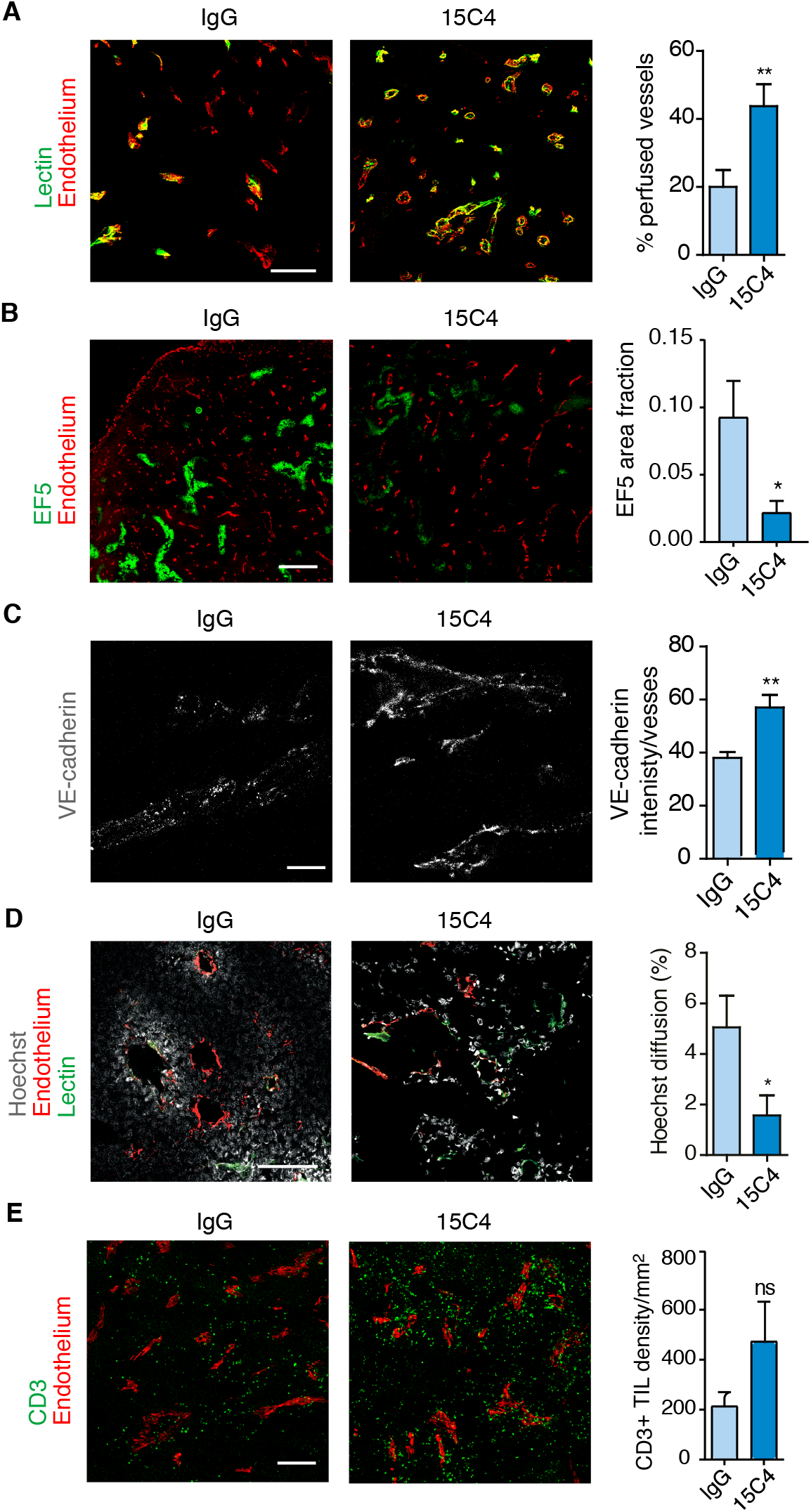
Loss or inhibition of LRG1 improves vascular function. Immunohistochemistry and quantification of (**A**) tumor vessel perfusion (lectin; *n*=12 tumors each condition; scale bar, 200 μm), (**B**) hypoxia (EF5; IgG, *n=*7; 15C4, *n*=5 tumors; scale bar, 1 mm), (**C**) adherens junction molecule (VE-cadherin; *n*=5 tumors each condition; scale bar, 50 μm), (**D**) permeability (Hoechst; IgG, *n*=11; 15C4, *n*=10 tumors; scale bar, 200 μm) and (**E**) tumor infiltrated lymphocyte density (CD3^+^ lymphocytes; IgG, *n*=13; 15C4, *n*=11 tumors; scale bar, 250 μm). For all graphs mean ± s.e.m. ns non-significant; Student t-test. *P<0.05, **P<0.01.

### Inhibition of LRG1 increases the delivery and efficacy of chemotherapy

The data above raised the possibility that vascular normalization, through inhibition of LRG1, may be exploited to enhance the delivery of additional therapeutic agents. To test whether LRG1 blockade enhances the efficacy of a co-therapy we first investigated 15C4 in combination with the cytotoxic agent cisplatin in the B16-F0 subcutaneous model. While both 15C4 and the maximum tolerated regimen of cisplatin each elicited a reduction in tumor size, their delivery in combination was significantly more effective (Fig. 6A and Supplementary Fig. 7). Analysis of the growth rates of individual tumors showed that the combined therapy of 15C4 and cisplatin was 25% more effective at inhibiting tumor growth than control IgG and cisplatin (Fig. 6B). Consistent with this, and the hypothesis that 15C4 enhances the delivery of cisplatin to the tumor, we saw a large increase in DNA double-stranded breaks, as revealed by *γ*-H2AX positivity (Fig. 6C), and apoptosis (Fig. 6D), demonstrating that inhibition of LRG1 improves the delivery, and hence efficacy, of a cytotoxic drug.

**Figure 6.**
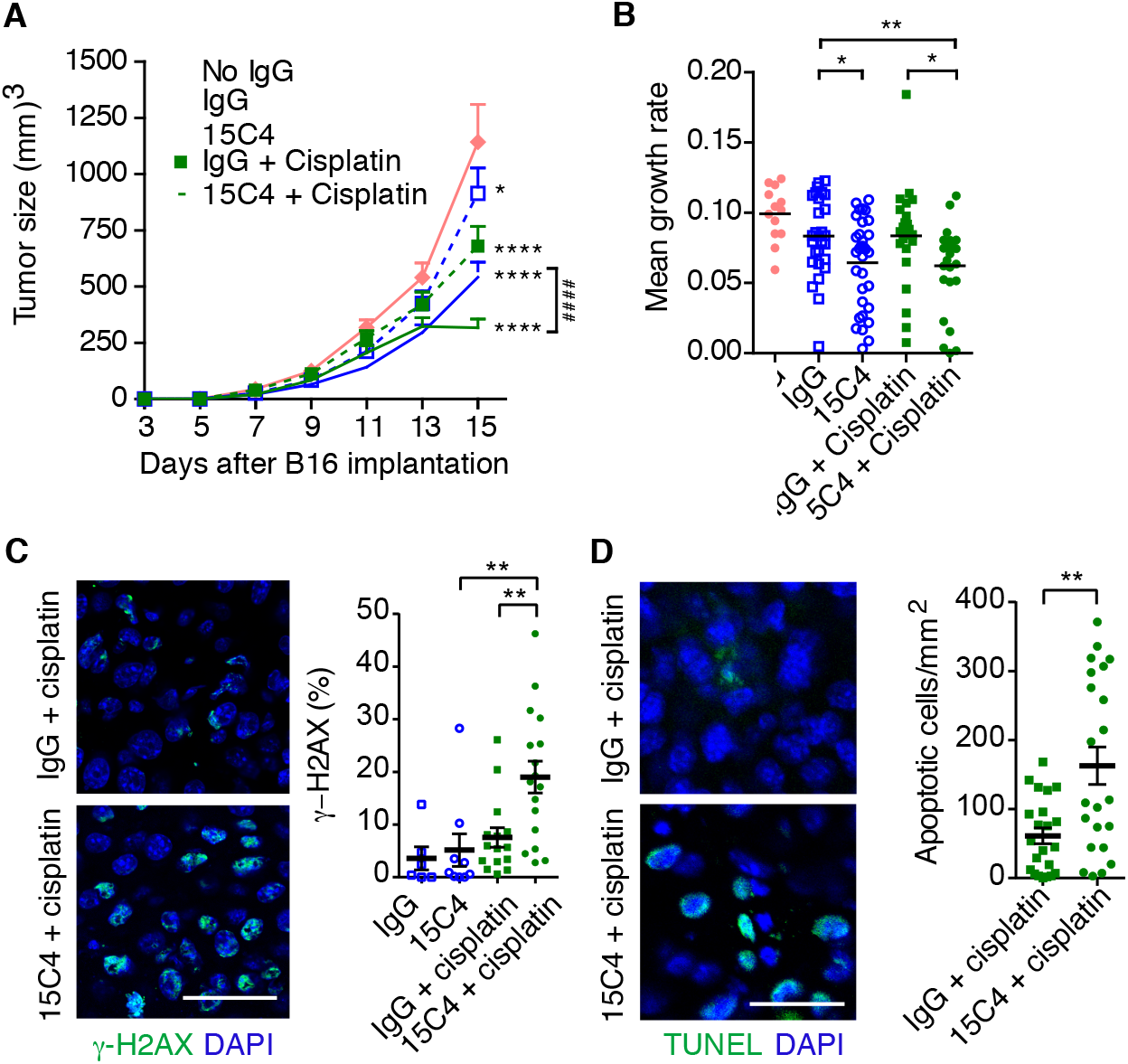
Antibody inhibition of LRG1 enhances the efficacy of cisplatin. Treatment of B16-F0 tumors with 15C4 (50 mg/kg) and cisplatin (2.5 mg/kg). (**A**) Growth curves (mean ± s.e.m.), analyzed by linear regression comparing to no IgG (*P<0.05, ****P<0.0001) or pairs as shown (^####^P<0.0001). No IgG, *n*= 13; IgG, *n*=28; 15C4, *n*=33; IgG + Cisplatin, *n*=23 and 15C4 + cisplatin, *n*=27 mice. (**B**) Growth rate (slope) of each tumor. Student t test, *P<0.05, **P<0.01. (**C**) DNA double strand breaks detected with antibody against γ-H2AX (green). Scale bar, 30μm. Graph shows percentage of γ-H2AX^+^ nuclei (mean ± 95% CI). One-way ANOVA, **P<0.01. IgG, *n*=6; 15C4, *n*=9; IgG + cisplatin, *n=15* and 15C4 + cisplatin, *n*=17 mice. (**D**) Apoptotic cells revealed by TUNEL staining (green). Graph shows density of TUNEL^+^ apoptotic cells (mean ± s.e.m.). Student t test, **P<0.01. IgG + Cisplatin, *n=22* and 15C4 + cisplatin, *n*=22 mice.

### LRG1 inhibition enhances the efficacy of adoptive cell therapy

These results led us to ask whether a similar enhancement of tumor cell killing could be achieved with other therapeutic modalities. In particular, we sought to establish whether in such a cold tumor we could enhance the effect of immunotherapies. We therefore investigated the combination of LRG1 antibody blockade and adoptive T cell therapy. Following subcutaneous grafting of B16-F10 melanoma cells harboring the internal influenza nucleoprotein antigen NP68 (NP68-B16) (*37*), donor F5B6 CD8^+^ T-cells expressing a TCR specific for the NP68 peptide were transferred to the tumor-bearing host mice. As previously described (*37*), F5B6 CD8^+^ T cells significantly reduced tumor growth (Fig. 7A and Supplementary Fig. 8). However, the combination of 15C4 with this dose of adoptive T cells led to a 30% greater reduction in tumor growth rate (Fig. 7A,B). Upon histological analysis the number of CD3^+^ T-cells that had infiltrated the tumor increased marginally in the 15C4 alone and donor CD8^+^ T cells groups but following combination therapy were elevated significantly, predominantly as a result of donor CD8^+^ T cell (CD90.2^+^) infiltration (Fig. 7C). The effects on tumor growth and T cell entry were replicated in a subsequent study in which the mice were treated with a reduced titer of F5B6 CD8^+^ T cells, but with the same dose of 15C4 antibody and extended for a further 13 days (Fig. 7D and Supplementary Fig. 9A). Again, an increase in tumor infiltrating-lymphocytes (TILs) was observed, particularly antigen-activated donor cells, which is consistent with vascular normalization and improved delivery. We also observed less peritumoral T cell cuffing and an associated increase in the density of intratumoral TILs (Supplementary Fig. 9B), suggesting that migration from the tumor vascular margin through the stroma is also enhanced in 15C4 treated tumors.

**Figure 7.**
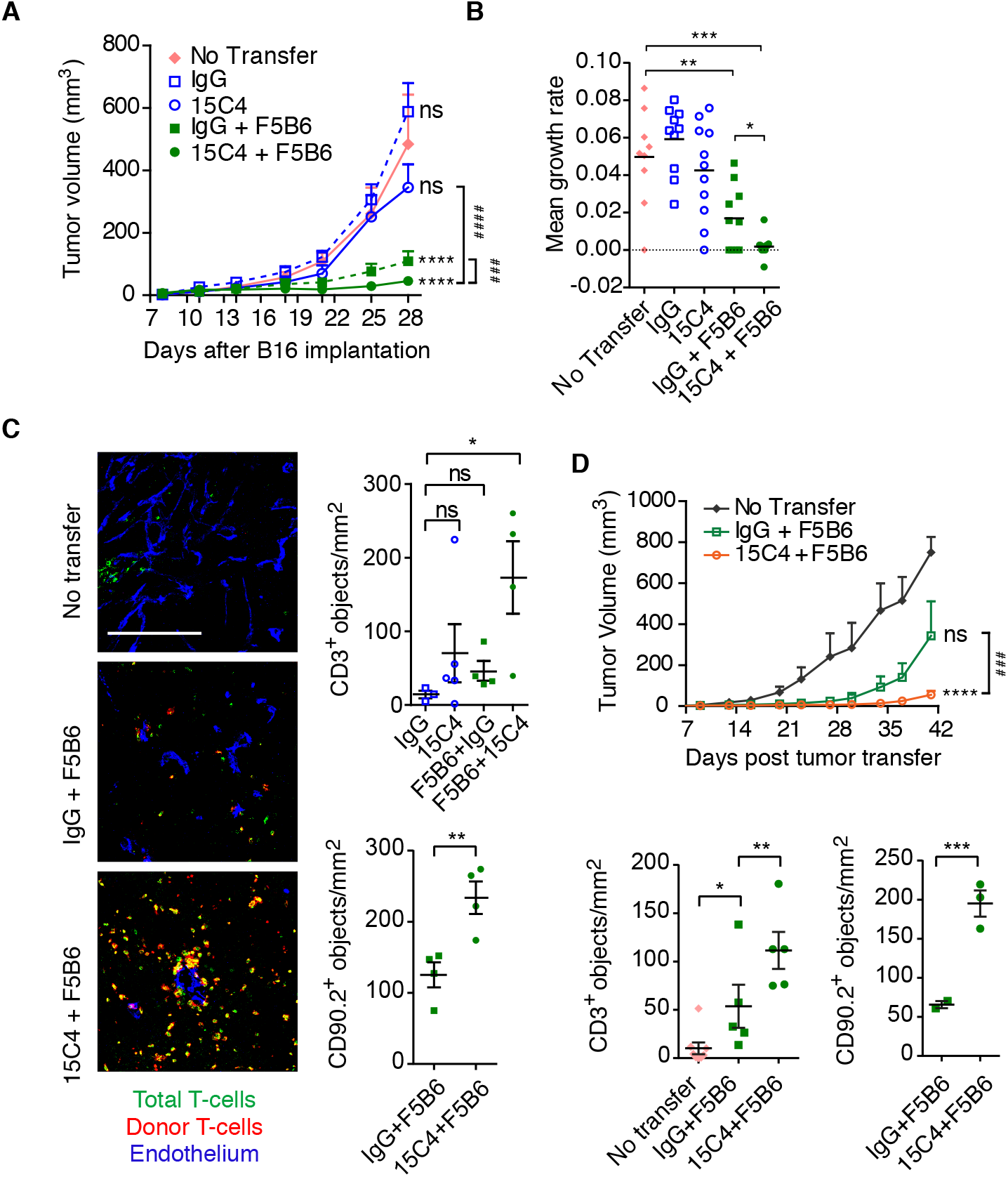
Antibody inhibition of LRG1 improves the efficacy of adoptive T cell therapy. Treatment of mice bearing NP68-expressing B16-F10 subcutaneous tumors with 15C4 and F5B6 cytotoxic T-cells. (**A**) Growth curves (mean ± s.e.m.), analyzed by linear regression comparing to No Transfer (****P<0.0001) or pairs as shown (^###^P<0.001, ^####^P<0.0001). No Transfer, *n*= 9; IgG, *n*=10; 15C4, *n*=11; IgG + F5B6, *n*=10 and 15C4 + F5B6, *n*=11 mice. (**B**) Growth rate (slope) of each tumor; Student t test, *P<0.05, **P<0.01, ***P<0.001. (**C**) T-cell infiltration of tumors. Scale bar, 200 μm. Graphs show density (objects/mm^2^) of CD3^+^ T-cells (top) and of CD90.2^+^ donor cells (bottom). Student t-test, *P<0.05, **P<0.01. (**D**) As in (**A**) but with reduced F5B6 cytotoxic T-cell treatment. Graphs show density (objects/mm^2^) of CD3^+^ T-cells (left) and of CD90.2^+^ donor cells (right). No transfer, *n*= 8; IgG + F5B6, *n*= 10; 15C4 + F5B6, *n*= 10; linear regression. *P<0.05, **P<0.01, ***P<0.001.

### LRG1 inhibition augments the effect of PD-1 checkpoint inhibition

The use of immune checkpoint antagonists, including inhibitors of CTLA-4, PD-1 and PD-L1, has proven to be very effective in the treatment of hematological cancers and melanoma but their impact on many solid cancers has been less effective. Having shown that 15C4 treatment enhanced the efficacy of adoptive T cell therapy we therefore investigated whether 15C4 could also augment the effectiveness of a checkpoint inhibitor. B16-F0 cells were grafted into wild type mice followed by treatment with 15C4, an anti-PD1 blocking antibody or a combination of both. As monotherapies, 15C4 and anti-PD1 each elicited a significant reduction in tumor volume (Fig. 8A. Supplementary Fig. 10) and mean growth rate (Fig 8B) with anti-PD-1 producing 33% tumor growth inhibition (TGI). In combination with 15C4, however, overall TGI was 88% with evidence of tumor regression occurring at the later time point. Histological analysis at study end point revealed increased cytotoxic CD8^+^ T cell infiltration in the combination therapy group (Fig. 8C,D). Consistent with this, we also observed enhanced granzyme B expression (Fig. 8E), indicative of greater cytotoxic lymphocyte activity. This enhancement of intratumoral immune activity is particularly striking in such an immunologically silent model.

**Figure 8.**
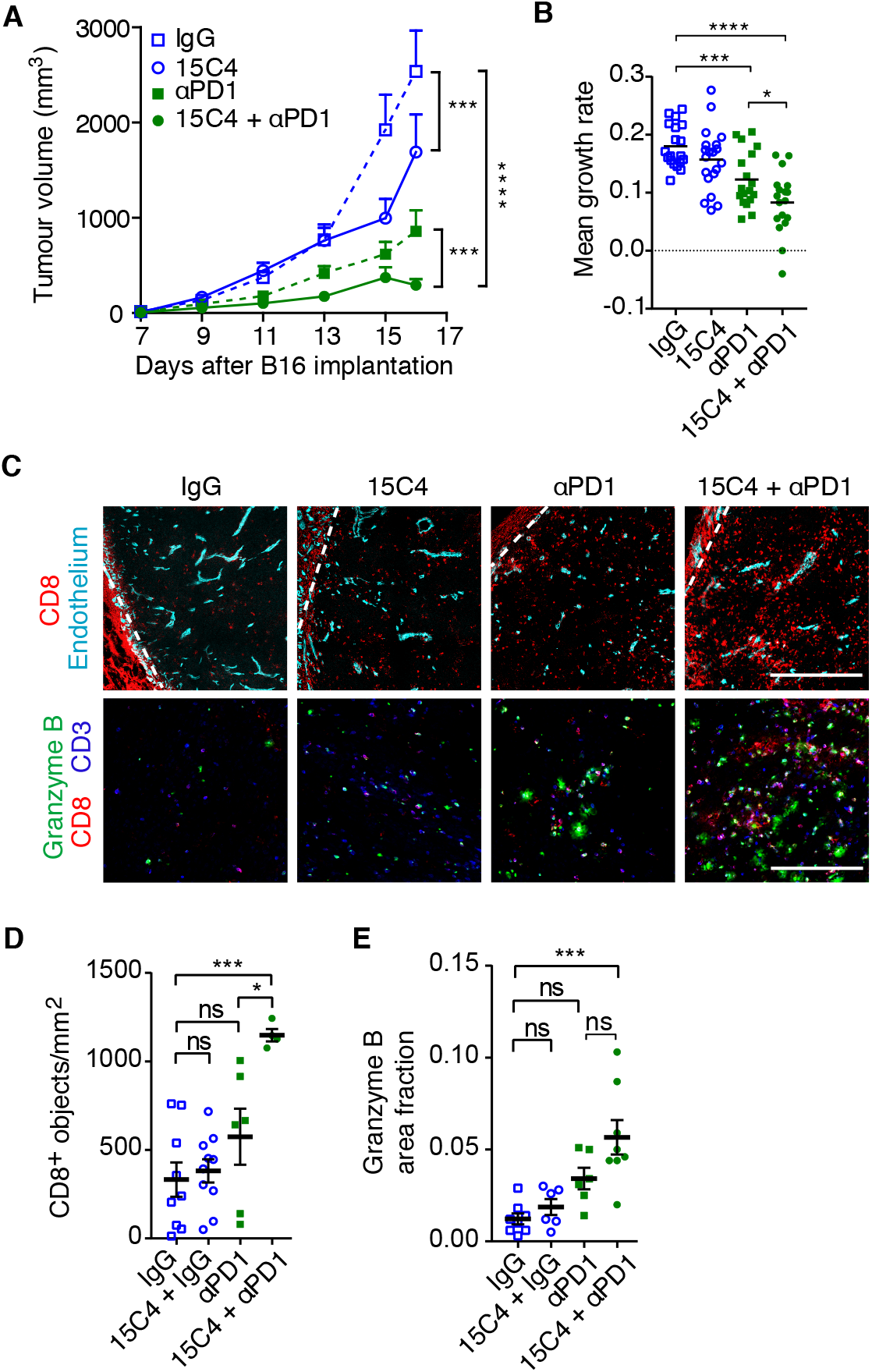
Antibody inhibition of LRG1 improves the efficacy of adoptive T cell therapy. Treatment of B16-F0 subcutaneous tumors with 15C4 and anti-PD-1. (**A**) Growth curves (mean ± s.e.m.), analyzed by linear regression comparing pairs as shown. (**B**) Growth rate (slope) of each tumor; Student t test. (**C**) T-cell infiltration of tumors. Dashed line represents tumor edge. Scale bars, 500 μm (top) and 200 μm (bottom). Graphs show density (objects/mm^2^) of CD8^+^ T-cells (**D**) and granzyme B area fraction (**E**). *n*= 9, 10, 6, 4 for **I** and *n*=8, 6, 6 and 8 for **M**, tumors, left to right; Student t-test, *P<0.05, **P<0.01, ***P<0.001.

### Combination of anti-LRG1 and anti-PD1 does not induce the formation of HEV

It has previously been shown that the combination of vascular normalizing agents and checkpoint inhibitors can stimulate the formation of HEV’s and that this may play a significant role in driving enhanced leukocyte recruitment and subsequent improved tumor cell killing (*38*). To determine whether our combined therapy also induced HEV formation we undertook qPCR analysis of a panel of HEV signature genes, namely glycosylation-dependent cell adhesion molecule 1 (*Glycam1*), and the chemokines *Ccl19*, *Ccl21* and *Cxcl13* (*14*). Expression of these genes was negligeable in the control and treatment arms with no significant differences between groups (Supplementary Fig 11) indicating no significant induction of HEVs. This was further confirmed by immunohistological staining of tumor sections with the MECA79 antibody, that detects peripheral node addressin (PNAd), revealing a lack of signal in control and treatment groups (Supplementary Fig. 11). These data indicate, therefore, that while HEV formation in other settings may be a contributing factor to the observed increase in TILs and treatment efficacy, it is not the only mechanism through which a combination of vascular normalizing strategies with immune checkpoint inhibition can elicit a beneficial effect.

## DISCUSSION

In this study we provide new insight into the cause of dysfunctional vessel growth in tumors and show that LRG1 is a significant angiopathic factor capable of disrupting the normal angiogenic process and contributing to the pro-oncogenic microenvironment of primary tumors. Under normal conditions the principal source of LRG1 is the liver where it may serve as an acute phase protein (*39*) involved in wound healing (*40*). In many diseases, however, LRG1 is induced locally in tissue lesions by the vascular endothelia and, in the case of cancer, by surrounding tumor cells. Locally produced LRG1 is known to contribute to the formation of abnormal vessels in the eye and kidney (*1,2*), in part by disrupting homeostatic TGFβ signaling in a highly context-dependent manner. Here we have shown that LRG1 also impacts on the vasculature of tumors and that deletion of the *Lrg1* gene, or inhibition of LRG1 function with a blocking antibody, improves vessel structure and function. Our data indicates that LRG1 is not directly pro-angiogenic but most likely facilitates neovascularization through its destabilizing effect on pericyte-endothelial cell interactions, a prerequisite for angiogenic sprouting (*41*). The decrease in vessel density observed in some models in the absence of *Lrg1* or following antibody blockade may therefore reflect vascular stabilization and subsequent suppression of angiogenesis. These improvements in vessel function were mediated independently of any alterations in gene expression of cardinal ligand-receptor pathways involved in determining vascular status, including members of the VEGF signaling network. One gene that was found to be altered significantly was the shear response factor *Klf2*. KLF2 is intimately involved in endothelial cell quiescence (*34*) and can be transcriptionally activated through TGFβALK5 signaling (*42*). This finding is consistent with LRG1 biasing endothelial cell TGFβ signaling in favor of the destabilizing ALK1-Smad 1/5/8 pathway (*1,2*) and away ALK5-Smad2/3 signaling. *Lrg1* deletion, therefore, enable maintenance of TGFβALK5 signaling and vascular quiescence.

The attendant decrease in tumor growth and improvement in survival with deletion or blockade of LRG1 on its own may be a consequence of decreased angiogenesis, or due to associated effects on the tumor microenvironment. For example, improved oxygenation resulting from better perfusion is likely to reverse hypoxia-mediated changes such as activation of pro-oncogenic HIF1 mediated genes, selection and expansion of aggressive clones and immune evasion (*43–46*). A further contributing factor may be a direct effect of LRG1 on cancer cells where it has been reported to stimulate their proliferation and migration (*47,48*). These deleterious effects of LRG1 are, however, at odds with a previous report of its suppression of LLC tumor growth (*49*), but are consistent with overwhelming clinical evidence that increased circulating LRG1 levels are diagnostic and associated with poor prognosis (Supplementary Table 1).

The discovery that LRG1 is angiopathic led us to test the hypothesis that blocking LRG1 would improve vascular function and augment the effects of other therapies. Treatment of tumor-bearing mice with 15C4 significantly enhanced the cytotoxic effect of cisplatin, suggesting that vascular normalization permits more effective delivery of the drug to the tumor mass. Indeed, combination of 15C4 and cisplatin exhibited not only reduced growth rate but also evidence of tumor regression that was not evident with monotherapy. The increased tumor cell death may also be enhanced through improved tumor oxygenation as hypoxia attenuates the effectiveness of cisplatin and contributes to chemotherapeutic resistance (*50*). Consistent with the cisplatin study, blockade of LRG1 also improved the efficacy of adoptive T cell therapy. The B16-F0 mouse tumor model is recognized as an immunologically cold tumor that, under normal conditions, exhibits very few TILs. This was confirmed in B16-F10 melanoma cells expressing the NP68 internal influenza nucleoprotein antigen as a surrogate tumor antigen, where very few infiltrated CD3^+^ T cells were seen in the untreated mice. Adoptive cell therapy with NP68 antigen-specific cytotoxic CD8^+^ T cells led to reduced tumor growth but this did not correlate with a significant increase in the number of infiltrated CD3^+^ T cells in end-stage tumors, as has been noted previously in this model (*51*). This is likely to reflect the dynamic nature of T cell involvement where single end-point analysis does not record possible temporal changes in T-cell recruitment, retention, exit, proliferation and death. Nevertheless, what was strikingly clear was that in the presence of LRG1 inhibition, adoptive T cell therapy led to greater tumor destruction and a significant increase in total CD3^+^ T cells that were predominantly donor cells. This undoubtedly reflects the antigen activated status of the donor cells and their enhanced migratory and retention capacity compared to circulating naïve T cells. The improved vascular patency brought about by blocking LRG1 therefore results in better access to the tumor vascular bed enabling greater penetration into the tumor mass (*52–54*). It is also likely that improved oxygenation will counteract some of the negative effects of hypoxia that may impact on T cell proliferation, retention and survival within the tumor microenvironment.

As with cisplatin and adoptive T cell therapy, blocking LRG1 with 15C4 also vastly improves the efficacy of a checkpoint inhibitor. It has long been recognized that tumors can evade immune rejection through eliciting powerful immunosuppressive signals that prevent an effective T cell response. This negative immune regulation can be impeded through the use of immune checkpoint inhibitors (*55,56*), which in certain human cancers has led to marked improvements in tumor destruction and overall survival. However, many cancers remain immunologically silent for reasons that are likely to be multifactorial (*57*), but which may include impaired immune cell delivery due to compromised perfusion (*53,54*). This has led to the concept that vascular normalization strategies targeting VEGF, or other vascular modulating factors such as angiopoietin 2, endothelial glycoprotein L1, notch 1 and regulator of G protein signaling 5 (Rgs5) (*22*), may enhance effector cell entry to the tumor. What such studies have revealed is a profound crosstalk between the vasculature and the tumor immune microenvironment. Accordingly, vascular normalization promotes the infiltration of immune cells but can also enhance the expression of immune modulators such as the checkpoint ligand PD-L1 (*14–16*). Consistent with this crosstalk it has been reported that abnormalization of the vasculature decreases immune cell infiltration and that deletion of CD4^+^ T cells promotes dysfunctional vessels (*25*) revealing the close functional interplay between the immune system and the vasculature.

Improved lymphocyte infiltration previously reported with vascular normalization has been attributed to the formation HEV (*38*). These structures, normally present in lymph nodes, are characterized by a plump morphology and expression of specialized adhesion molecules, including peripheral node addressin (PNAd), that facilitate leukocyte traffic. Their induction in tumors, therefore, has provided a mechanistic explanation for the increased presence of TILs and enhanced tumor killing observed in combination therapies. In our setting, however, we were unable to observe the presence of HEV as determined by a panel of distinguishing markers. Other factors, over and above improved perfusion, may therefore be responsible for the enhanced infiltration observed. One feature of tumor endothelial cells that may contribute to poor leukocyte recruitment is their failure to respond effectively to activation by inflammatory mediators, a condition termed endothelial cell anergy (58). This manifests in part as loss of expression of key cell adhesion molecules, such as ICAM-1 and VCAM-1 (*59,60*), which are required for effective leukocyte recruitment (*61*). Here we show that vascular normalization, brought about by *Lrg1* deletion, reverses in part this anergy through enhancing the induction of *Icam1* and *Vcam1* which in turn will facilitate the recruitment of circulating leukocytes (*62*). This demonstrates that in addition to HEV formation, other mechanisms brought about by vascular normalization are at play in facilitating leukocyte tumor infiltration.

Our finding that inhibiting LRG1 with a function-blocking antibody reverses its detrimental effects on the tumor vasculature and enhances both adoptive T cell and immune checkpoint inhibition strategies lends further weight to the view that improving vascular function is a promising co-therapeutic strategy. Targeting LRG1, therefore, may provide an additional, or alternative, approach for normalizing the tumor vasculature and enhancing the efficacy of co-therapies. At present the principle approach is to employ anti-VEGF axis inhibitors which have shown some capacity to normalize the vasculature and improve immunotherapies (*14*). However, there remain major challenges with this approach not least of which is the difficulty in determining the appropriate dose and the purported short therapeutic window (*6,9*). This is confounded by difficulties in determining the relative activity of the VEGF axis in different tumors (*13*). Unlike VEGF targeted therapies, blocking LRG1 has the potential advantage that patients may be stratified as higher circulating levels generally correlate with a worse prognosis (Supplementary Table 1).

As interest in vascular normalization increases, various targets other than those of the VEGF axis have been identified (*22*) but for most their clinical utility remains untested. Here we present LRG1 as a promising target but its safety and successful translation into patients need to be proven. Unlike some targets, however, *Lrg1* knockout in the mouse does not produce an overt phenotype and they remain fertile and healthy over a normal lifespan providing evidence that it is not critical to homeostasis. In addition, as an ectopic non-essential modifier of TGFβ signaling, LRG1 blockade may offer advantages over direct therapeutic targeting of the TGFβ superfamily for treating cancer, which in general has been disappointing. Failure in this area is likely due to the difficulty in separating homeostatic from pathogenic TGFβ signaling as many core TGFβ signaling components are involved in both. TGFβ operates as an analogue signaling network whose effects are largely determined by a balance of complex, and nuanced, interactions between different arms of the extensive signaling cascade. Under normal conditions homeostatic TGFβ signaling is required for a stable vasculature but during disease this is disturbed and LRG1 is a prime disrupting candidate. However, targeting endoglin, which is upregulated on neovascular endothelia and which is a binding partner of LRG1, has proven to be disappointing in achieving a therapeutic effect on tumor angiogenesis, although any impact on vascular normalization has not been fully investigated. This failure may be due, in part, to antagonizing a binding site important for maintaining vascular quiescence (*63*). Interestingly, LRG1 binding to ENG facilitates the reconfiguration of the TFGβ receptor complex to enhance pathogenic signaling but in so doing may also result in loss of beneficial homeostatic BMP9/ENG signaling. Targeting LRG1, therefore, removes an independent pathogenic factor that disturbs the homeostatic balance in TGFβ signaling without interfering with essential components of the network.

In conclusion, we have shown that LRG1 is a major driver of abnormal vessel growth in solid primary tumors and that its inhibition leads to significant restoration of normal vascular function. This raises the possibility that therapeutic targeting of LRG1 will improve the quality of vessels not only in cancer, but in diseases as diverse as diabetic kidney disease, neovascular age-related macular degeneration, and inflammatory disease, and pave the way towards improved strategies to revascularize ischemic tissue.

## MATERIALS AND METHODS

### Cell Culture

Cancer cell lines B16-F0 (mouse melanoma) and LLC1 (LL/2; mouse Lewis Lung carcinoma) were maintained in Dulbecco’s Modified Eagle’s Medium supplemented with glucose (4.5 g/L), sodium pyruvate (110 mg/L), 10% FCS, penicillin (100,000 U/L) and streptomycin sulphate (100 mg/L). Cultures were maintained at 37^0^C in 5% CO_2_ and checked to be clear of mycoplasma contamination.

### Tumor models

All procedures were performed in accordance with the UK Animals (Scientific Procedures) Act and the Animal Welfare and the Ethical Review Bodies of the UCL Institute of Ophthalmology, Cancer Research UK Beatson Institute, University of Glasgow, and Cardiff University.

#### Subcutaneous graft models

C57BL/6 mice were purchased from Harlan Laboratories. *Lrg1*^−/−^ mice were generated by the University of California Davies knockout mouse project (KOMP) repository (http://www.komp.org/). Single-cell suspensions of 1×10^6^ B16-F0 or LLC cells were injected subcutaneously into the lower back of *Lrg1^+/+^* or *Lrg1*^−/−^ male C57BL/6 mice in 100 μl PBS. Mice were randomized by age prior to inoculation. Tumors were measured without bias at defined intervals using calipers and tumor volume was calculated using the formula: V = (4/3) × π × (L/2) × (W/2) × (H/2). Mice were sacrificed at the end of the experiment, or when tumors exceeded 1000 mm^3^. The mean tumor growth rate for individual tumors was calculated using the slope of log transformed tumor volumes (*56*).

#### Genetically engineered mouse models

Mice were housed in the animal facility at the CRUK Beatson Institute. All experiments were performed on a C57BL/6 background. For ageing experiments of spontaneous models (*Apc^Min/+^* and KPC) mice were aged and sampled when showing moderate signs of illness. Tumors in *villin^CreER^ Apc^fl/+^* mice were induced, at an age of 6-10 weeks by a single intra peritoneal injection of 2 mg Tamoxifen (Sigma; T5648) in corn oil (Sigma; C8267) and aged until showing moderate signs of illness. No distinction between males and females has been made in all mouse experiments and blinded for *Lrg1* status.

### Immunohistochemistry

Subcutaneous B16-F0 and LLC tumor models and KPC tumors were fresh frozen on dry-ice and embedded in optimal-cutting-temperature medium (OCT). Contiguous frozen tissue sections were cut at a thickness of 8 μm and stored at −20°C. Sections were fixed in 4% paraformaldehyde, 100% methanol or 100% acetone, depending on antibodies used. The small intestine and colon from *Apc^Min/+^* and the *vil^CreER^ Apc^fl/+^* tumors were formalin-fixed, paraffin-embedded (FFPE). 5 μm sections were deparaffinized and immunolabelled following antigen retrieval. In all cases sections were blocked in 0.5% BSA and washed in 0.01% tween-20 in PBS. Antibodies used to label mouse endothelium were anti-CD31 (Dianova or Abcam), endomucin (Abcam), VE-cadherin (insert) or podocalyxin (R&D systems) was used in B16-F0 tumors, as it strongly labelled the endothelium and the staining pattern was almost indistinguishable from CD31 in this model (Supplementary Fig. 4). Pericytes were labelled with antibodies to NG2 (Merck-Millipore) or αSMA (Sigma-Aldrich). Antibodies to basement membrane proteins collagen-IV (Merck-Millipore) or perlecan (Abcam), and immune cell markers CD3 (Abcam), CD8 (Novus), or CD90.2 (Biolegend) were also used. Other primary antibodies were to granzymeB (Novus) and EF5 (Merck-Millipore). Alexa-fluor labelled secondary antibodies were from Thermofisher.

### RNAScope^®^ *in situ* hybridisation

FFPE tumor or intestine samples were placed in xylene followed by absolute ethanol. For chromogenic detection, slides were processed using the 2.0 HD Detection kit – BROWN (Advanced Cell Diagnostics) and the manufacturer’s instructions. For fluorescent detection, slides were processed using the Multiplex Fluorescent Kit v2, followed by TSA^®^ signal amplification (PerkinElmer), and immunohistochemistry performed afterwards if desired. Slides were hybridized with probes specific to *Lrg1* and quality of signal and tissue determined using positive (*Ppib*) and negative (*Dapb*) probes, supplied by the manufacturer (Advanced Cell Diagnostics). The specificity of the *Lrg1* probe was confirmed by probing tumors sections from *Lrg1*^+/+^ *and Lrg1^−/−^* mice (Supplementary Fig. 11).

### Analysis of vessel density and normalisation

To measure vessel profiles in the tumors, tumor sections were labelled with antibodies to tumor endothelium markers (CD31, PDXL or endomucin). B16F0 and LLC sections were imaged using a Nikon Eclipse Ti epifluorescence microscope (Nikon). The entire tumor vasculature was included in the analysis, excluding vasculature in the tumor periphery. KPC sections were imaged on a Zeiss 700 confocal microscope. At least two 850×850 μm areas per section containing tumor vessels were imaged and maximum intensity projections of z-stacks analyzed. Vessel density (number per unit area) and size (cross-sectional area) were calculated from thresholded images from B16-F0, LLC and KPC tumors using NIS-Elements software (Nikon). Vessels were identified as objects between 5-800 μm^2^ that were positive for the endothelial marker. The mean vessel size and density per tumor section is reported. For *Apc^Min/+^* and the *vil^CreER^ Apc^fl/+^* sections, at least 2 intestinal adenomas per mouse were imaged, using a Zeiss 700 confocal microscope and the mean result reported. Vessels in a 250×188 μm ROI at the luminal edge of the adenoma were analyzed. Since vessels were mostly contiguous in these images, vessel area per image was calculated, using ImageJ, rather than vessel size and density of individual vessels.

The association of pericytes or basement membrane proteins with the tumor endothelium was measured from sections labelled with antibodies to endothelial cells (CD31, endomucin or podocalyxin) and multiple pericyte (NG2 and/or αSMA) or matrix (perlecan and collagen IV) markers. For pericytes, a 0.37 or 0.72 cm^2^ ROI encompassing the edge and core of the tumor was imaged and then analyzed using NIS elements software (Nikon) or ImageJ (http://imagej.nih.gov/ij/). For analysis of endothelial basement membrane, at least 2 640×640 μm areas per section containing tumor vessels were imaged on a Zeiss 710 microscope and maximum intensity projections of z-stacks analyzed. The fraction of perlecan or collagen IV pixels which overlap CD31 positive pixels was calculated from a single plane though the center of the z-stack. The same threshold was used for each image and Manders' overlap coefficient was calculated using JACoP plugin on ImageJ. Data was normalized to the mean control value for each experiment. Images were blinded before analysis in all cases.

### Tumor hypoxia and vascular perfusion

To measure tumor hypoxia, 0.2 ml of 10mM EF5 (Merck-Millipore) was injected into the peritoneum of tumor bearing mice and tumors harvested after 1 hr. Pimonidazole adducts in sections were detected by immunohistochemistry using anti-EF5, clone ELK3-51 Cyanine 3 conjugate and the entire tumor section imaged using a Nikon Eclipse Ti epifluorescence microscope. The proportion of each tumor positive for hypoxia stain was measured from identically thresholded images on NIS elements software (Nikon) and reported as a percentage of total image area.

To examine tumor vessel perfusion and leakage, tumor bearing mice were injected intravenously with FITC-labelled Lycopersicon esculentum lectin (Vector labs; 10 mg/kg) and low molecular weight fluorescent DNA binding dye Hoechst 33342 (Sigma-Aldrich; 7.5 mg/kg), respectively, followed 3 min later by perfusion fixation with 4% paraformaldehyde. Cryosections were labelled with an antibody to endomucin to count endothelialized vessels. The percentage of perfused vessels was calculated as the % of endomucin-positive vessels which were also lectin positive. The proportion of each ROI positive for Hoechst was measured from thresholded images on NIS elements software (Nikon), and normalized to lectin area, i.e. perfused vessels.

### Tumor co-therapy strategies

#### Chemotherapy

To investigate the effect of LRG1 blockade on efficacy of tumor chemotherapy, wild-type C57BL/6 mice were injected with B16-F0 cells subcutaneously into the flank and treated with 50 mg/kg of the function-blocking anti-Lrg1 monoclonal antibody 15C4 or IgG control (rat or mouse IgG1, Supplementary data, Fig 10B) administered by intraperitoneal injection every 3 days from day 3. At day 7, a maximum tolerated dose (2.5 mg/kg) of the chemotherapy drug cisplatin were administered every other day by intraperitoneal injection until the mice were sacrificed at the end of the experiment or when tumors exceeded 1000 mm^3^. Cisplatin-induced DNA damage was assayed using an antibody against the DNA double strand break marker gamma-H2AX (Merck-Millipore) on tumor sections co-stained with DAPI to enumerate cell nuclei. The percentage of nuclei with gamma-H2AX foci was measured from confocal images (Zeiss 700) which were blinded before analysis. Apoptotic cells were identified by TUNEL assay on sections using an ApopTag in situ apoptosis detection kit (Merck-Millipore).

#### Adoptive T cell therapy

To investigate the effect of LRG1 blockade on efficacy of tumor immunotherapy a mouse model of adoptive T cell therapy was used as described (Watson et al, 2016). Briefly, 5 × 10^5^ NP68-B16 melanoma cells in 200μl sterile PBS were injected subcutaneously into the shaven left flank of B6.PL-Thy1a/CyJ (Thy1.1/CD90.1) or C57BL/6 (Thy1.2/CD90.2) mice, tumors grown for 6 days and the mice sub-lethally irradiated with 597cGy total body irradiation. On day 7, F5B6 CD8^+^ T cells (> 95% naive (CD62L positive, CD44 low) CD8^+^ T cells) expressing the F5 T cell receptor for NP68 peptide on a C57BL/6 background were isolated from spleens of naïve F5B6 mice using a CD8α+ T cell isolation kit for negative selection, and LS columns, according to the manufacturer’s instructions (StemCell Technologies). Briefly, spleens were harvested from adult mice and mashed through a 70 μm cell strainer (BD Pharmingen). Red blood cells were lysed using red cell lysis buffer (Biolegend) and lymphocytes washed with ice-cold phosphate buffered saline (PBS) supplemented with 2 % fetal calf serum (FCS) prior to magnetic isolation. The enriched CD8+ cell fraction was counted using a hemocytometer, resuspended in sterile PBS for injection and analyzed for CD8, CD62L, CD44, CD27 and F5 TCR expression.

Tumor bearing mice were randomly distributed into 5 treatment groups of 8-11 mice (No transfer; IgG; 15C4: IgG + F5B6; 15C4 + F5B6) and injected subcutaneously with peptide vaccine (100μg NP68 peptide in 200μl incomplete Freund’s adjuvant) into the right flank prior followed by 2.25 × 10^5^ F5B6 CD8^+^ T cells (CD90.2) injected into the tail vein. Mice were treated with 50 mg/kg of the function-blocking anti-Lrg1 mouse monoclonal antibody 15C4 or IgG control administered by intraperitoneal injection commencing on the same day as T cell transfers and antibody administration repeated every 3 days until the end of the study. Tumors were measured with calipers at defined intervals and tumor volume was calculated using the formula: V = (4/3) × π × (L/2) × (W/2) × (H/2). At the end of the experiment, mice were sacrificed, blood collected for serum and tumors fresh frozen on dry-ice in optimal-cutting-temperature medium (OCT) and stored at −80°C before immunostaining tumor infiltrating T cells either for total T cells (CD3^+^) or for donor T cells (CD90.2 in tumors grown in CD90.1 mice).

#### Immune checkpoint blockade

To investigate the effect of LRG1 blockade on efficacy of PD-1/PD-L1 axis blockade in an immunologically cold tumor, wild-type C57BL/6 mice were injected with 1 × 10^6^ B16-F0 cells subcutaneously into the flank. Mice were treated with a combination of 50 mg/kg of the function-blocking anti-LRG1 15C4, 200 μg rat anti-mouse PD1 (Bio X Cell) or 200 μg rat IgG2a isotype control (Bio X Cell). Mice were dosed by intraperitoneal injection commencing on day 3 and antibody administration was repeated every 3 days until the end of the study. Tumors were measured with calipers at defined intervals and tumor volume was calculated. At the end of the experiment, mice were sacrificed and tumors were fresh frozen on dry-ice in OCT and stored at −80°C before immunostaining.

### T-cell infiltration analysis

Fresh-frozen sections were fixed in 100% ice-cold methanol and/or 4% formaldehyde and labelled using antibodies to total T cells (CD3^+^), donor T-cells (CD90.2^+^), cytotoxic T cells (CD8^+^) and/or granzyme B. For each section a 2920×2920 μm or 4250×4250 μm tile scan was acquired encompassing the edge and core of the tumor, using Zeiss 710 confocal microscope. Maximum intensity projections of z-stacks were analyzed using NIS elements software (Nikon). CD3^+^, CD8^+^ and CD90.2^+^ objects were identified by thresholding and automatically enumerated. Granzyme B^+^ area from 4250×4250 μm images was identified by thresholding and reported as a fraction of total image area.

### Scanning Electron Microscopy

14 days after subcutaneous B16F0 injection *Lrg1^+/+^* and *Lrg1*^−/−^ mice bearing tumors were perfusion fixed in Karnovsky fixative (2% paraformaldehyde, 2.5% Glutaraldehyde), followed by immersion fixation in fixative overnight at 4° C. Vibratome sections (200 μm) were washed in PBS and then osmicated with 1% osmium tetroxide in ddH_2_O for 1 hour. They were then washed in ddH_2_O and dehydrated in alcohol. The tumor sections were then immersed in dry methanol and in hexamethyldisilazane (reagent grade >99%, Aldrich chemicals) and then allowed to dry. The specimens were fixed onto aluminum stubs using a conductive carbon disc and silver paint (Agar) and were then coated with 2 nm platinum in a Cressington sputter coater. Imaging was done on a Zeiss Sigma VP SEM using the in lens detector.

### RNASeq and RT-qPCR

RNA from *Lrg1^+/+^* and *Lrg1*^−/−^ B16F0 tumors was extracted using the RNeasy mini kit (Qiagen) and analyzed for quality using the 4200 TapeStation (Agilent). mRNA was prepared from total RNA for sequencing using the Kapa riboerase library preparation kit (Agilent) and was sequenced for differential expression analysis (0.5 High output NextSeq run, 43bp paired end reads). Raw RNA sequence data has been deposited with NCBI Sequence Read Archive accession number PRJNA552723 (https://www.ncbi.nlm.nih.gov/sra/PRJNA552723). For RT-qPCR analysis, total RNA was isolated from B16-F0 tumors that were treated with 15C4/PD1 as indicated in the experimental conditions. cDNA was synthesised using the LunaScript RT SuperMix Kit (New England Biolabs E3010) and gene expression was analyzed by RT-qPCR on QuantStudio 6 (Applied Biosciences) using the Luna Universal qPCR kit (New England Biolabs, M3003). Relative expression was normalized to *Actb* and *Gapdh* housekeeping genes and was determined using the ΔΔCt method. Primer sequences for the mouse genes were as follows: *Ccl19*, Forward: CAGTCACTCCCCTGTGAACC, Reverse: CAGAGTTGGGGCTGGGAAG, *Ccl21a*, Forward: AAGGCAGTGATGGAGGGGGT, Reverse: CTTAGAGTGCTTCCGGGGTG, *Cxcl13*, Forward: CAGGCCACGGTATTCTGGA, Reverse: CAGGGGGCGTAACTTGAATC, *Glycam1*, Forward: TCAGCTGCAACCACCTCAG, Reverse: TTCGTGATACGACTGGCACC.

### Statistical analysis

Images were blinded before analysis in all cases. Statistical analysis was performed using Graphpad Prism version 5.0 or 7.0 for Windows, GraphPad Software, La Jolla California USA, www.graphpad.com. Error bars and statistical tests used for each experiment are indicated in the figure legends. All t tests were two-tailed. A *P<* value less than 0.05 was considered statistically significant. Grubb’s test was used to test for outliers (www.graphpad.com).

## LIST OF SUPPLEMENTARY MATERIALS

**Supplementary data Figure 1**. Normal colon, pancreas and skin do not express *Lrg1*.

**Supplementary data Figure 2**. B16-F0 and LLC tumor growth in individual *Lrg1*^−/−^ and *Lrg1*^+/+^ mice.

**Supplementary data Figure 3**. Immunohistochemical staining for tumor vascular density and association with basement membrane in colorectal cancer models.

**Supplementary data Figure 4**. Expression of key genes involved in vascular maturation or destabilization.

**Supplementary data Figure 5**. Gene expression of key endothelial cell adhesion molecules involved in leukocyte recruitment.

**Supplementary data Figure 6**. Effect of antibody blockade of LRG1 on B16-F0 tumor growth and vasculature.

**Supplementary data Figure 7**. Individual B16-F0 tumor growth rates in mice treated with 15C4, cisplatin or 15C4 plus cisplatin.

**Supplementary data Figure 8**. Individual B16-F0 tumor growth rates in mice treated with 15C4, adoptive T cells or 15C4 plus adoptive T cells.

**Supplementary data Figure 9**. Individual tumor growth rates and immunohistochemical detection of T cell infiltrates in mice treated with 15C4 and adoptive T cells.

**Supplementary data Figure 10**. Individual B16-F0 tumor growth rates from mice treated with 15C4, anti-PD-1 or a combination of both.

**Supplementary data Figure 11**. Expression of HEV markers in B16-F0 tumors.

**Supplementary data Figure 12**. Validation of *Lrg1* in situ hybridization and control antibodies.

**Supplementary Table 1**. Human cancer studies indicating LRG1 as a potential diagnostic and prognostic biomarker

## ACKNOWLEDGEMENTS

JG and SM would like to thank Jack Blackburn for his contribution towards 15C4 antibody production. We would like to thank the Biological Resources Unit at the UCL Institute of Ophthalmology and the Joint Biological Services at Cardiff University. We also thank Core Services and Advanced Technologies at the Cancer Research UK Beatson Institute (C596/A17196), with particular thanks to the Biological Services Unit and Histology.

## Author contributions

JG and SEM conceived, designed and directed the overall study. AA designed the adoptive T cell experiments and OJS and RJ the genetically engineered mouse model experiments. XW undertook the initial B16F0 experiment. MO’C and DMK performed all subsequent experimental work using the B16F0 and LL2 models. RJ and OJS conducted the experimental work using the genetic cancer models. MO’C and DMK undertook all the histological staining and analysis. AHW, MA and JO conducted the adoptive T cell experiments. MO’C, DMK, CC, CP and AD performed the analysis of immune cell infiltration into tumours. LD and AD undertook the gene analysis studies. JGe purified and characterised the 15C4 blocking antibody.

## Data and materials availability

All data from the main text and supplementary materials is available.

