## Supplementary figures and images for "LRG1 destabilizes tumor vessels and restricts immunotherapeutic potency"

### Supplemental Figure 1

#### Supplementary Figure 1

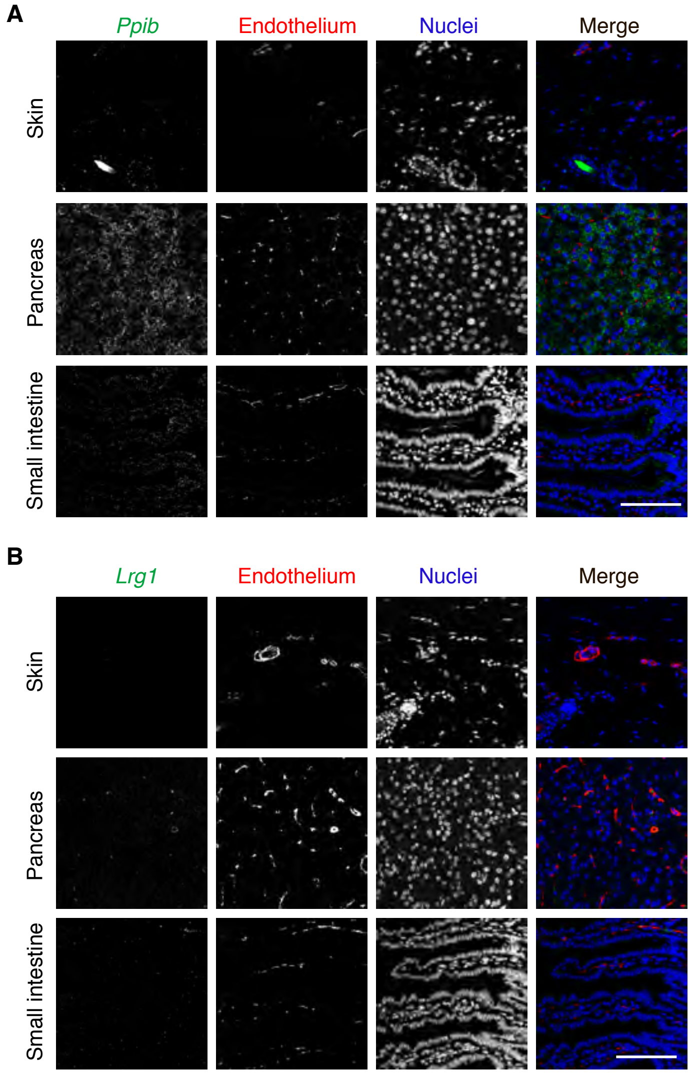

### Supplemental Figure 2

##### Supplementary Figure 2

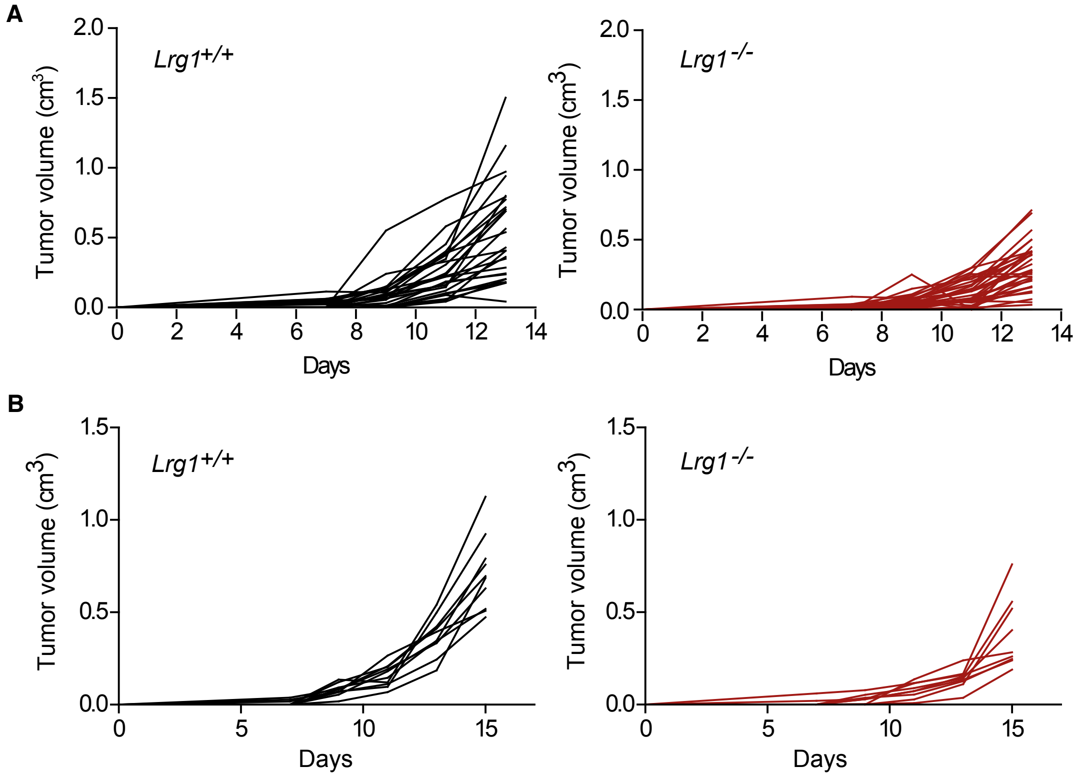

### Supplemental Figure 3

##### Supplementary Figure 3

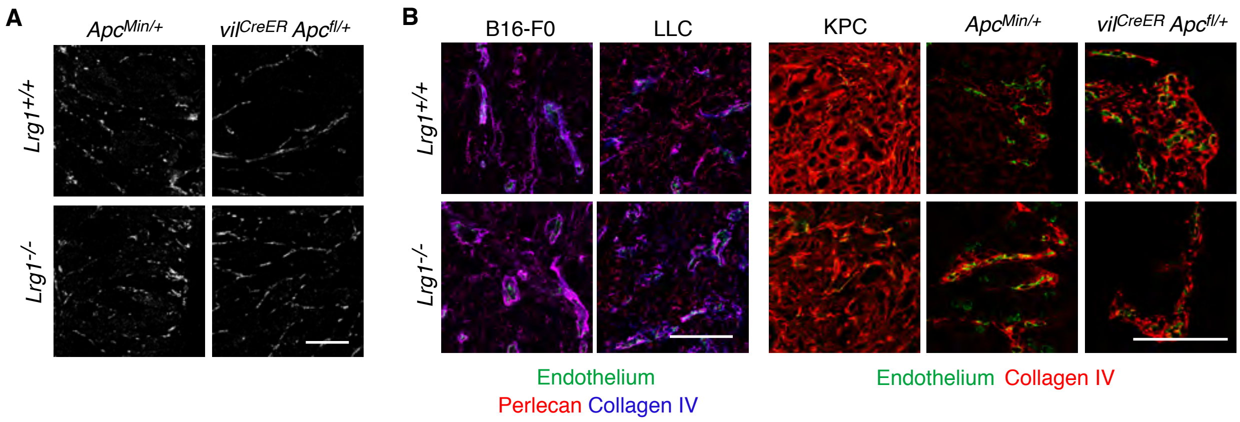

### Supplemental Figure 4

##### Supplementary Figure 4

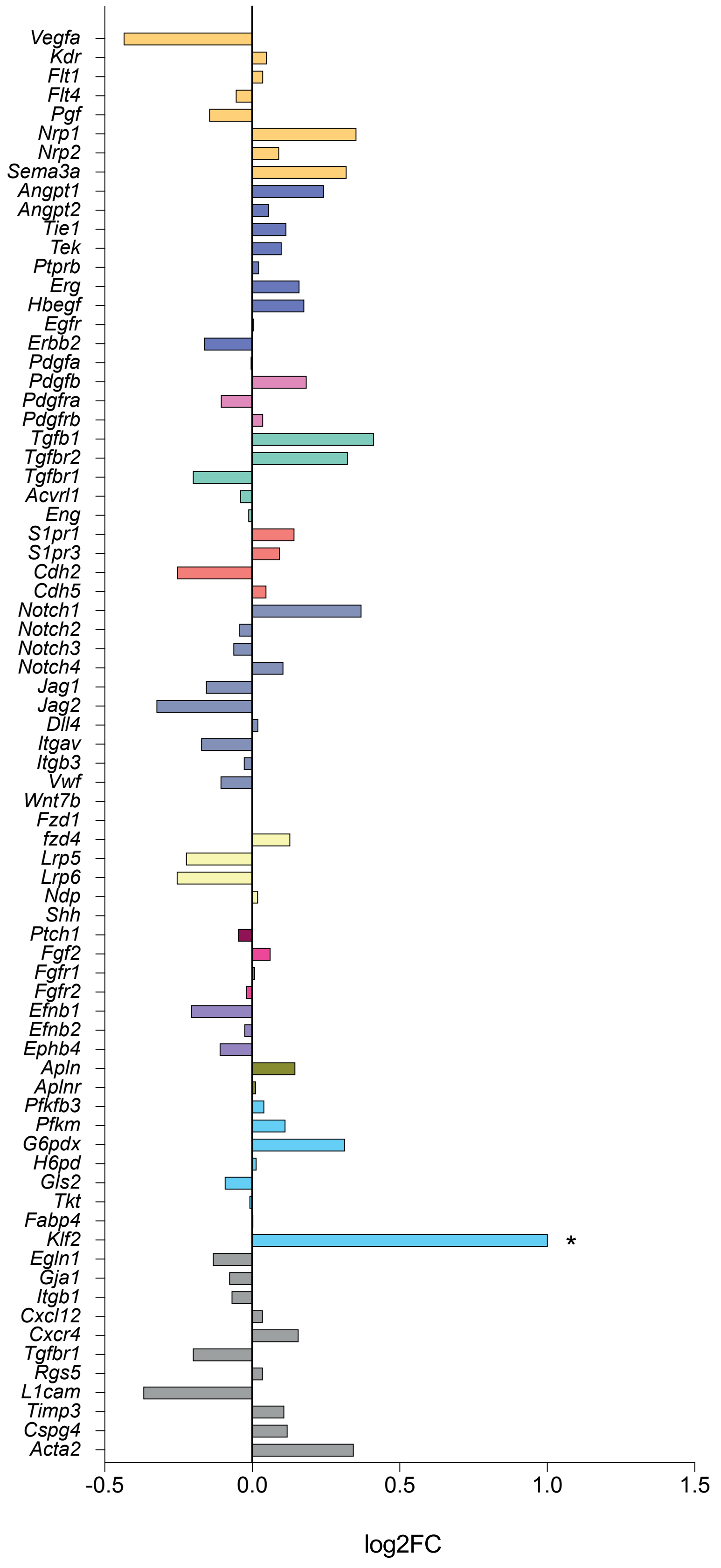

### Supplemental Figure 5

**Supplementary Figure 5**

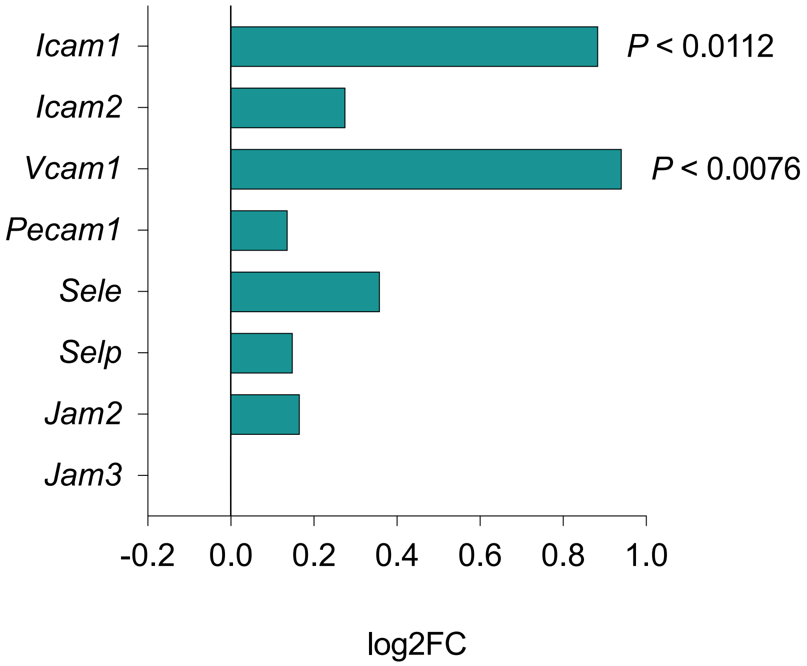

### Supplemental Figure 6

##### Supplementary Figure 6

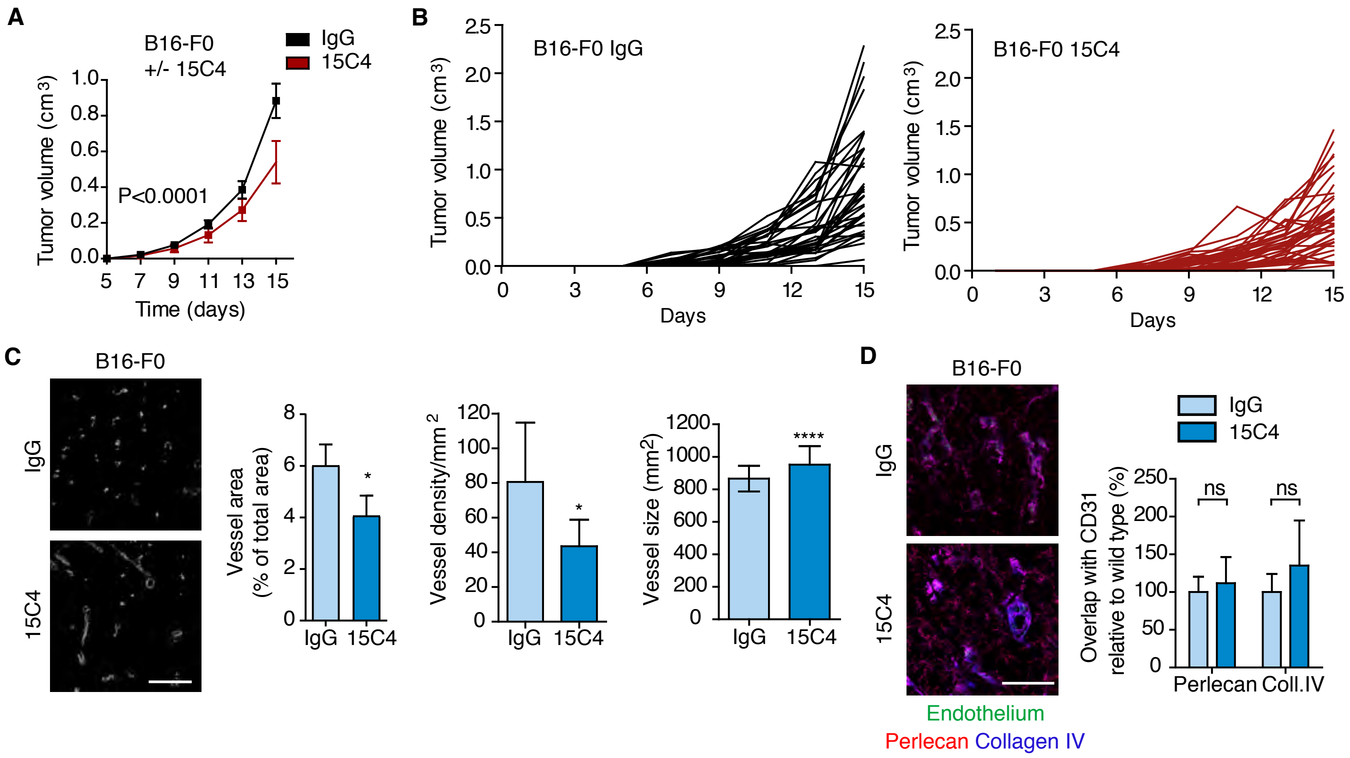

### Supplemental Figure 7

##### Supplementary Figure 7

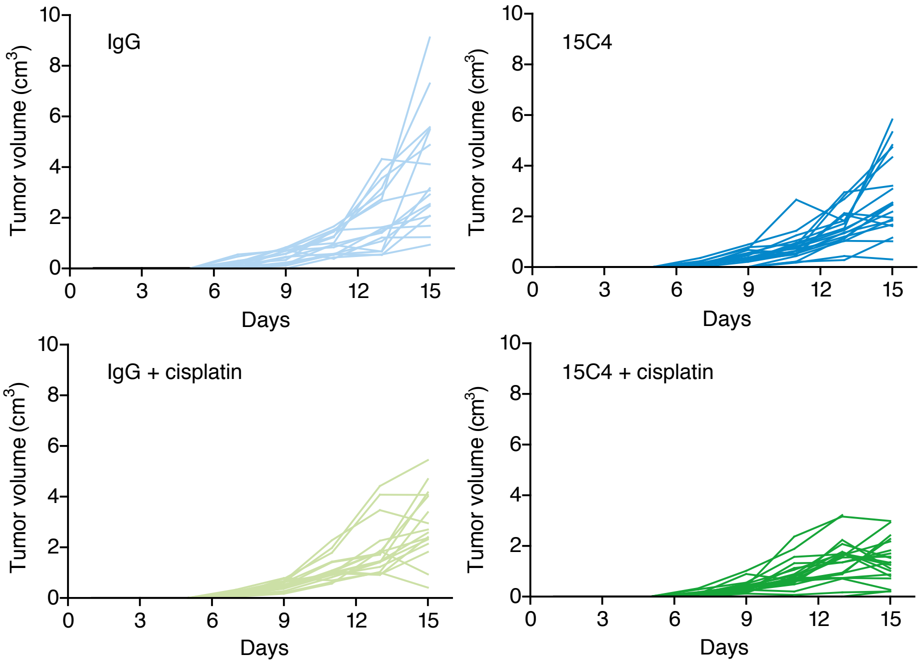

### Supplemental Figure 8

##### Supplementary Figure 8

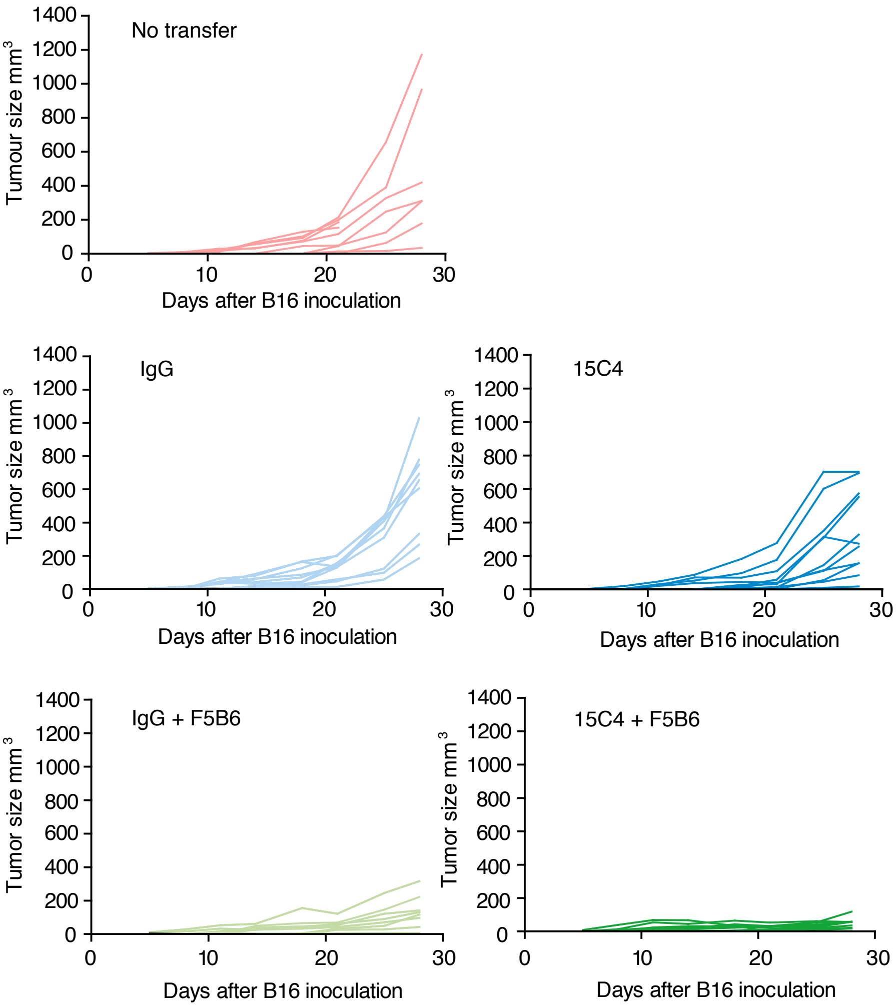

### Supplemental Figure 9

#### Supplementary Figure 9

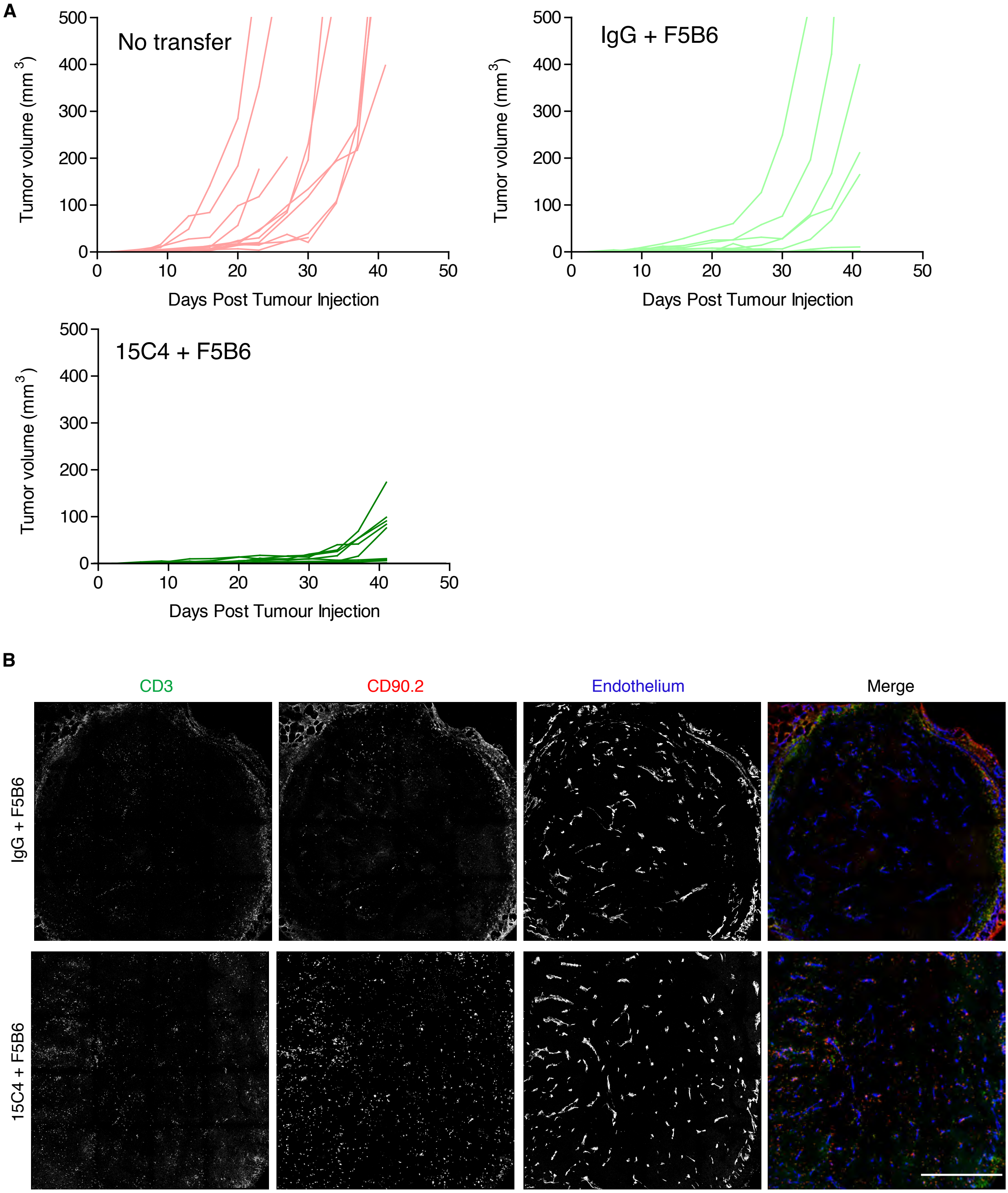

### Supplemental Figure 10

##### Supplementary Figure 10

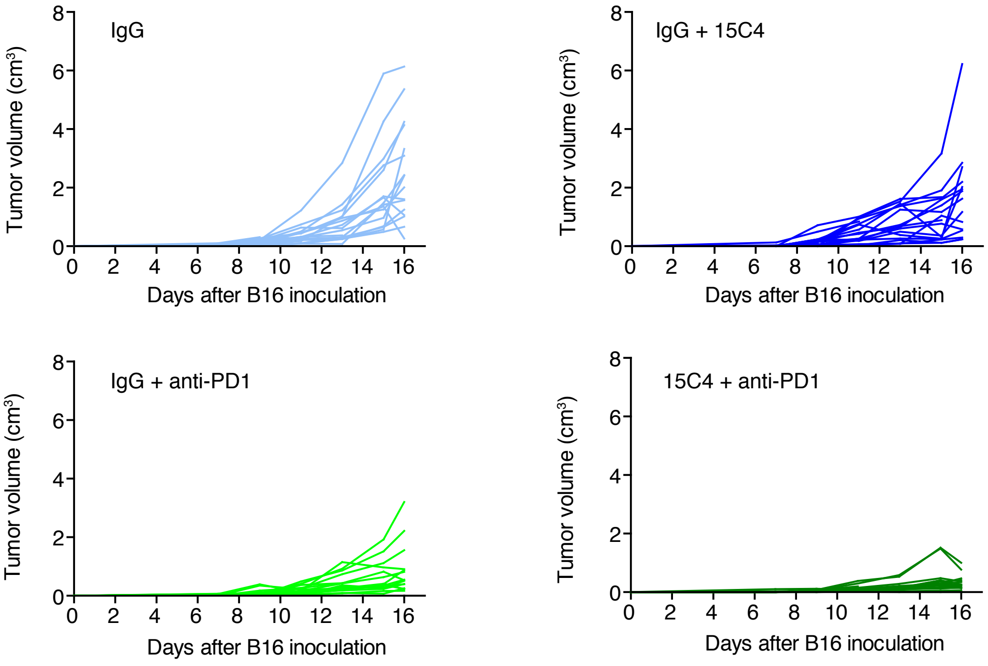

### Supplemental Figure 11

##### Supplementary Figure 11

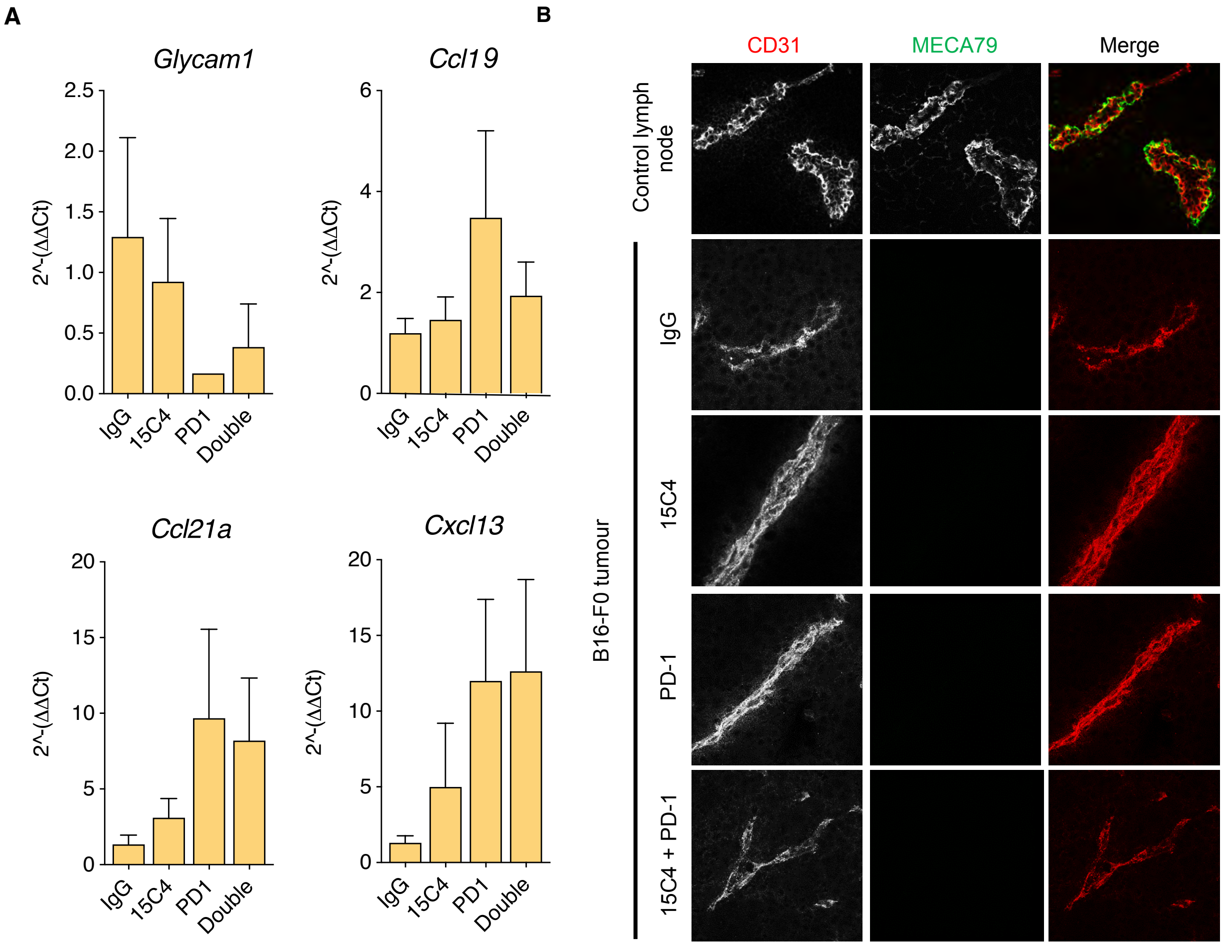

### Supplemental Figure 12

##### Supplementary Figure 12

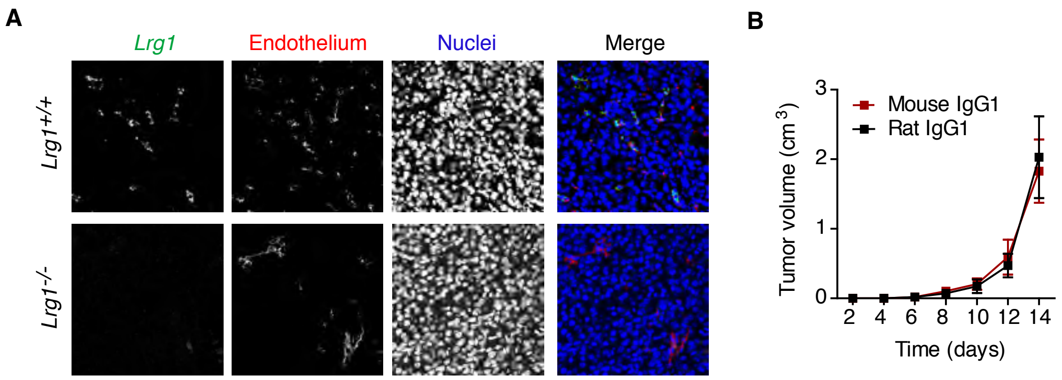
