## Supplementary Data Figure legends for "LRG1 destabilizes tumor vessels and restricts immunotherapeutic potency"

**Supplementary data Figure 1. Normal colon, pancreas and skin do not express *Lrg1*.** RNAScope® in situ hybridization of sections from wild-type (non-tumor bearing) mouse tissue using probes to positive control transcript *Ppib* (**A**) or to *Lrg1* (**B**) followed by immunohistochemical staining of endothelial cells (CD31). Scale bar, 100 μm.

**Supplementary data Figure 2. B16-F0 and LLC tumor growth in individual *Lrg1*$^{-/-}$ and *Lrg1*$^{+/+}$ mice.** Effect of *Lrg1* deletion on individual B16-F0 (**A**) and LLC (**B**) subcutaneous tumor volume over time.

**Supplementary data Figure 3. Immunohistochemical staining for tumor vascular density and association with basement membrane in colorectal cancer models.** (**A**) CD31 (PECAM-1) stained sections of the vasculature from *Apc*$^{Min/+}$ and *vil*$^{CreER}$ *Apc*$^{fl/+}$ adenomas from *Lrg1*$^{+/+}$ and *Lrg1*$^{-/-}$ mice. Scale bar, 50 μm (**B**) Endothelial basement membrane association with tumor vessels in sections labelled with antibodies to CD31, perlecan and/or collagen IV. Scale bar, 100 μm.

**Supplementary data Figure 4. Expression of key genes involved in vascular maturation or destabilization.** Expression of indicated genes, grouped into families, in B16-F0 tumors from *Lrg1*$^{-/-}$ mice compared to wild type controls. RNASeq data expressed as fold change. * $P < 0.05$.

**Supplementary data Figure 5. Gene expression of key endothelial cell adhesion molecules involved in leukocyte recruitment.** Expression of indicated genes in B16-F0 tumors from *Lrg1*$^{-/-}$ mice compared to wild type controls. RNASeq data expressed as fold change.

**Supplementary data Figure 6. Effect of antibody blockade of LRG1 on B16-F0 tumor growth and vasculature.** (**A**) Mean tumor volumes of B16-F0 tumors from wild-type mice treated with anti-LRG1 (15C4) or control antibody (IgG) (mean ± 95% CI). IgG, *n*=35 mice; 15C4, *n*=39 mice; RM two-way ANOVA. (**B**) Individual growth curves of tumors from wild-type mice treated with anti-LRG1 (15C4) or control antibody (IgG). (**C**) Vascularity of tumors revealed by CD31 immunohistochemistry. Scale bar, 250 µm. Graphs show % $CD31^+$ area in the image, and vessel density and cross-sectional area of individual $CD31^+$ objects (mean ± 95% CI). IgG, *n*=14 tumors; 15C4, *n*=18 tumors; Mann Whitney test. (**D**) Endothelial basement membrane association with tumor vessels from wild-type mice treated with IgG or 15C4. Sections were labelled with antibodies to CD31, perlecan and/or collagen IV. Scale bar, 100 µm. Overlap with endothelium was measured and normalized to mean control value in each experiment (mean ± sem). IgG *n*=12, 15C4 *n*=13; Mann Whitney test, ns non-significant. *P<0.05, ****P<0.0001.

**Supplementary data Figure 7. Individual B16-F0 tumor growth rates in mice treated with 15C4, cisplatin or 15C4 plus cisplatin.** Individual growth curves of tumors from different treatment arms of experiment shown in Fig. 3A.

**Supplementary data Figure 8. Individual B16-F0 tumor growth rates in mice treated with 15C4, adoptive T cells or 15C4 plus adoptive T cells.** Individual growth curves (from Fig. 7A) of tumors from mice bearing NP68-expressing B16-F10 subcutaneous tumors treated with 15C4 and F5B6 cytotoxic T-cells.

**Supplementary data Figure 9. Individual tumor growth rates and immunohistochemical detection of T cell infiltrates in mice treated with 15C4 and adoptive T cells.** (**A**) Individual growth curves of tumors (from Fig 7D) from mice bearing NP68-expressing B16-F10 subcutaneous tumors treated with 15C4 and F5B6 cytotoxic T-cells. (**B**) Immunohistochemical detection of $CD3^+$ and donor T cell (CD90.2) infiltration into NP68-expressing B16-F10 subcutaneous tumors from mice treated with adoptive T cells alone or a combination of adoptive T cells and 15C4. Scale bar, 1 mm.

**Supplementary data Figure 10. Individual B16-F0 tumor growth rates from mice treated with 15C4, anti-PD-1 or a combination of both.** Individual growth curves of tumors (from Fig. 8A) from mice bearing B16-F0 subcutaneous tumors and treated with 15C4 and anti-PD-1.

**Supplementary data Figure 11. Expression of HEV markers in B16-F0 tumors. (A)** qPCR analysis of HEV signature genes in B16-F0 tumors from mice treated with control IgG, 15C4, anti-PD-1 antibody and a combination of 15C4 and anti-PD-1. (**B**) Immunohistochemical staining of lymph node and B16-F0 tumors with an anti CD31 antibody and the MECA79 antibody to detect PNAd. Tumors were taken from mice treated with control IgG, 15C4, anti-PD-1 antibody and a combination of 15C4 and anti-PD-1.

**Supplementary data Figure 12. Validation of *Lrg1* in situ hybridization and control antibodies.** (**A**) *Lrg1* RNAScope® probe is specific for mouse *Lrg1* transcript. Tumor sections from *Lrg1*$^{+/+}$ and *Lrg1*$^{-/-}$ mouse tumors, processed using *Lrg1* RNAScope® probe *in situ* hybridization followed by immunohistochemistry using antibodies to endothelial cells (CD31) and pericytes (αSMA). Scale bar, 100 μm. (**B**) Growth curves of B16-F0 tumors from wild-type mice treated with rat or mouse IgG1 (mean ± 95% CI). RM two-way ANOVA. *n*=8 mice for each treatment.
