## Supplementary Table 1 for "LRG1 destabilizes tumor vessels and restricts immunotherapeutic potency"

**Table 1. Human cancer studies indicating LRG1 as a potential diagnostic and prognostic biomarker**

| Cancer type | Detected | Conclusions | Reference link |
|---|---|---|---|
| **Biliary tract** | Serum | In conjunction with other biomarkers significantly elevated in serum in biliary tract carcinoma, particularly in high-risk patients with primary sclerosing cholangitis. | https://pubmed.ncbi.nlm.nih.gov/21970875/ |
| | Tumor tissue | Upregulation in tumour tissue providing a candidate independent prognostic factor. | https://pubmed.ncbi.nlm.nih.gov/32278969/ |
| **Bladder** | Urine | Significantly elevated with other markers. Proposed as diagnostic and for monitoring of recurrence. | https://pubmed.ncbi.nlm.nih.gov/22065568/ |
| **Breast** | Tumor tissue | LRG1 mRNA increased and potential biomarker for neo-adjuvant aromatase inhibitor treatment. | https://pubmed.ncbi.nlm.nih.gov/27491861/ |
| **Cervical** | Urine | Highly expressed in cervical cancer and diagnostic with other biomarkers | https://pubmed.ncbi.nlm.nih.gov/31186765/ |
| **Colorectal** | Plasma | Significantly elevated and with other predictive biomarkers proposed as a diagnostic and prognostic factor. Correlates with switch from adenoma to carcinoma and size of tumor. | https://pubmed.ncbi.nlm.nih.gov/22277732/ https://pubmed.ncbi.nlm.nih.gov/26253080/ https://pubmed.ncbi.nlm.nih.gov/26253081/ https://pubmed.ncbi.nlm.nih.gov/28697446/ https://pubmed.ncbi.nlm.nih.gov/29785123/ |
| | Serum | Significantly elevated and with other biomarkers diagnostic and prognostic. Associated with altered LRG1 glycosylation. | https://pubmed.ncbi.nlm.nih.gov/30971492/ https://pubmed.ncbi.nlm.nih.gov/31586890/ https://pubmed.ncbi.nlm.nih.gov/31753726/ https://pubmed.ncbi.nlm.nih.gov/29642865/ |
| | Tumor tissue | Overexpressed and associated with cancer aggressiveness and vascular density. Proposed as a diagnostic marker in general and prognostic marker for stage III colorectal cancer. | https://pubmed.ncbi.nlm.nih.gov/26856989/ https://pubmed.ncbi.nlm.nih.gov/29029535/ https://pubmed.ncbi.nlm.nih.gov/29785123/ |
| | Stool | With panel of biomarkers for early detection of high-risk adenomas and colorectal cancer. | https://pubmed.ncbi.nlm.nih.gov/31784980/ |
| **Endometrial** | Tumor tissue | Increased expression and independent prognostic factor of stage and lymphatic metastasis | https://pubmed.ncbi.nlm.nih.gov/24760273/ |
| **Gastric** | Serum and tumor tissue | Increased in serum and tissue. Proposed prognostic factor. | https://pubmed.ncbi.nlm.nih.gov/28746773/ |
| **Glioblastoma** | Plasma | With other markers showed positive correlation with tumor size | https://pubmed.ncbi.nlm.nih.gov/29513714/ |
| | Tumor tissue | Significantly higher than in lower-grade glioma. Proposed as potential diagnostic, prognostic, and regional biomarker. | https://pubmed.ncbi.nlm.nih.gov/32647893/ |
| **Hepatocellular** | Serum | Significantly elevated as part of a panel of protein biomarkers and associated with poor responders following transarterial chemoembolization | https://pubmed.ncbi.nlm.nih.gov/24195504/ https://pubmed.ncbi.nlm.nih.gov/28112944/ |

**Table 1. Human cancer studies indicating LRG1 as a potential diagnostic and prognostic biomarker**

| | | | |
|---|---|---|---|
| | Tumor tissue | Significantly up-regulated and prognostic for tumor size, differentiation, stage and vascularity. | https://pubmed.ncbi.nlm.nih.gov/26517349/ |
| **Lung** | Plasma | Significantly elevated in patients with non-small cell lung carcinoma and in panel of other biomarkers proposed as diagnostic. Highly indicative of reduced survival time post radiotherapy. | https://pubmed.ncbi.nlm.nih.gov/21947365/ https://pubmed.ncbi.nlm.nih.gov/26425690/ |
| | Serum | Significantly elevated and with other markers prognostic in patients with squamous cell lung carcinoma | https://pubmed.ncbi.nlm.nih.gov/23284758/ https://pubmed.ncbi.nlm.nih.gov/23042802/ |
| | Urine | Candidate biomarker for diagnosis of non-small cell lung carcinoma | https://pubmed.ncbi.nlm.nih.gov/21557262/ |
| | Tumor tissue | Upregulated in non-small cell lung carcinoma | https://pubmed.ncbi.nlm.nih.gov/23284758/ https://pubmed.ncbi.nlm.nih.gov/31528707/ |
| **Esophageal** | Plasma | Significantly elevated in esophageal squamous cell carcinoma and with alpha-2-HS-glycoprotein is a potential biomarker for early diagnosis. | https://pubmed.ncbi.nlm.nih.gov/25973038/ |
| | Serum | Increased and in combination with CRP and sIL-6R promising biomarker candidates to predict response to preoperative chemoradiotherapy | https://pubmed.ncbi.nlm.nih.gov/31582665/ |
| | Tumor tissue | Up-regulated and closely correlated with worse clinical survival | https://pubmed.ncbi.nlm.nih.gov/30714525/ |
| **Oral** | Plasma | Increased and with apolipoprotein A-IV are biomarkers for oral cancer screening and early diagnosis | https://pubmed.ncbi.nlm.nih.gov/30987719/ |
| | Serum | Increased in oral squamous cell carcinoma and with other biomarkers be an early diagnostic tool | https://pubmed.ncbi.nlm.nih.gov/25272005/ |
| | Saliva | Significantly elevated in oral squamous cell carcinoma and with other biomarkers associated with increased risk. | https://pubmed.ncbi.nlm.nih.gov/26552850/ |
| **Ovarian** | Serum | Significantly elevated and alone or with other biomarkers promising diagnostic factor. | https://pubmed.ncbi.nlm.nih.gov/20162585/ https://pubmed.ncbi.nlm.nih.gov/20831812/ https://pubmed.ncbi.nlm.nih.gov/20546617/ https://pubmed.ncbi.nlm.nih.gov/23731285/ https://pubmed.ncbi.nlm.nih.gov/25799488/ |
| | Urine | Significantly elevated and proposed biomarker. | https://pubmed.ncbi.nlm.nih.gov/24982608/ |
| **Pancreatic** | Plasma | Significantly elevated and may have utility with other markers as a diagnostic tool. Elevated during formation of intraductal papillary mucinous neoplasm and potential detection of early stage disease. | https://pubmed.ncbi.nlm.nih.gov/17303479/ https://pubmed.ncbi.nlm.nih.gov/26561977/ https://pubmed.ncbi.nlm.nih.gov/28376157/ https://pubmed.ncbi.nlm.nih.gov/29190982/ https://pubmed.ncbi.nlm.nih.gov/30575211/ https://pubmed.ncbi.nlm.nih.gov/30137376/ https://pubmed.ncbi.nlm.nih.gov/32545216/ |
| | Serum | Significantly elevated and with CA19-9 potential diagnostic biomarker. LRG1 glycosylation pattern changes significantly. | https://pubmed.ncbi.nlm.nih.gov/25058884/ https://pubmed.ncbi.nlm.nih.gov/16970316/ |

**Table 1. Human cancer studies indicating LRG1 as a potential diagnostic and prognostic biomarker**

| | | | |
|---|---|---|---|
| | Tumor tissue | Associated with higher recurrence rate and worse recurrence-free survival | https://pubmed.ncbi.nlm.nih.gov/30575211/ |
| **Renal** | Tumor tissue | Overexpressed in clear cell renal cell carcinoma and negatively related to patient survival | https://pubmed.ncbi.nlm.nih.gov/32337221/ |
| **Retinal** | Tumor tissue | Protein highly expressed in retinoblastoma | https://pubmed.ncbi.nlm.nih.gov/29392314/ |
